# A bipolar disorder-associated ultra-rare *AKAP11* protein-truncating variant attenuates stimulus-dependent PKA activation and induces anxiety- and depression-related behaviors in mice

**DOI:** 10.64898/2026.09.19.752887

**Authors:** Anna X. Luo, Leon Deng, Yannan Li, Xiaodi Zhang, Junnan Li, Xiaolei Zhu, Xiaobo Mao, Yijing Su, Bin Wu, Christopher A. Ross, Pan P. Li

## Abstract

Rare genetic variants associated with psychiatric disorders are thought to have larger individual effect sizes and direct biological roles, offering a unique opportunity for understanding causal relationships between genetic architecture and disease phenotypes. A recent whole-exome sequencing meta-analysis identified an increased burden of ultra-rare protein-truncating variants (PTVs) in *AKAP11* in patients with bipolar disorder and schizophrenia. *AKAP11* encodes an A-kinase anchoring protein that mediates the association of protein kinase A (PKA) and its substrates. How ultra-rare *AKAP11* variants contribute to psychiatric disorder pathogenesis remains unknown. Here, using CRISPR-Cas9 genome editing, we introduced into the mouse *Akap11* locus an ultra-rare PTV identified in a human patient with bipolar disorder. This ultra-rare PTV reduced *Akap11* mRNA and protein levels without producing stable novel mRNA isoforms, leading to *Akap11* haploinsufficiency. Male heterozygous mice exhibited behavioral abnormalities consistent with anxiety- and depression-related phenotypes. Transcriptomic and proteomic profiling of the neocortex revealed molecular changes involving synaptic organization, synaptic signaling, and neurite morphogenesis. Notably, the heterozygous mutant showed selective elevation in type I, but not type II, PKA regulatory subunits, as well as PKA catalytic subunits, in the neocortex. Functionally, these changes were accompanied by reduced phosphorylation of PKA substrates, attenuated stimulus-dependent PKA activity in cortical excitatory neurons, and impaired nascent dendritic development in cortical neurons. Together, our study presents a clinically relevant and genetically precise mouse model that provides mechanistic insight into how ultra-rare PTVs may contribute to psychiatric disorder pathogenesis, and a platform for testing rational therapeutics.

## Introduction

Psychiatric disorders possess complex genetic architectures characterized by high levels of polygenicity and pleiotropy [1, 2]. Genome-wide association studies (GWAS) have identified common genetic variants (minor allele frequency [MAF] greater than 1%) associated with psychiatric disorders [3–5], each contributing a small effect [6, 7]. More recently, whole-exome sequencing (WES) and whole-genome sequencing (WGS) studies have revealed rare variants (MAF < 1%) and ultra-rare variants (MAF < 0.1%) contributing to complex neurobiological disorders, including schizophrenia [8–10], bipolar disorder [11], autism spectrum disorder [12] and developmental disorder [13]. While rare variants explain a smaller fraction of population-level heritability, they often have larger individual effects and more direct biological roles [2, 6], offering unique opportunities to understand causal relationships between genetic architecture and disease phenotypes.

Bipolar disorder and schizophrenia affect 35.7 million and 26.9 million people worldwide, respectively, posing substantial healthcare burden [14]. A recent WES meta-analysis identified enrichment of ultra-rare protein-truncating variants (PTVs) in *AKAP11* in patients with bipolar disorder and schizophrenia [11]. *AKAP11* has also been predicted as a target gene of schizophrenia-associated *trans*-expression quantitative trait loci (*trans*-eQTLs) [15].

*AKAP11* encodes an A-kinase anchoring protein (AKAP) that mediates the association of protein kinase A (PKA) and its substrates [16, 17]. Prior *Akap11* loss-of-function mouse models generated by exon deletion (hereafter referred to as knockout [KO] models) showed altered PKA subunit expression, dysregulated striatal PKA dynamics, disrupted electroencephalogram signals, and anxiety- and depression-related behavioral deficits [18–21]. However, deletion of large genomic segments of *Akap11*, including exon 7 and/or exon 6, may disrupt *cis*-regulatory elements or local chromatin architecture. For example, exon 7 contains predicted distal enhancer-like elements and chromatin-accessible elements with CTCF binding signals [22].

Thus, exon-deletion models may have unexpected effects beyond disruption of the *Akap11* coding sequence.

In contrast, the effects of ultra-rare PTVs on *AKAP11* expression, downstream molecular pathways, and the disease-associated phenotypes remain largely unexplored. Moreover, a mouse model with a patient-derived ultra-rare PTV in the *Akap11* coding sequence would provide a more precise genetic perturbation than the exon-deletion models, and allow investigation of how rare coding variants contribute to bipolar disorder and schizophrenia pathogenesis.

Here using the CRISPR-Cas9 genome editing approach, we introduced a single-nucleotide variant to mouse *Akap11* exon 7 homologous to an ultra-rare PTV in human *AKAP11* exon 8 (ENST00000025301) identified in a bipolar disorder patient [11]. Exon 8 of human *AKAP11* not only contains binding sites for PKA, protein phosphatase 1, and GSK3β [16, 17], but also has the largest number (13 out of 16) of PTVs associated with bipolar disorder in the Bipolar Exome (BipEx) database [11]. The selected variant, chr13:42299894C>A, corresponds to c.1148C>A in the *AKAP11* coding sequence, and is predicted to convert Ser383 into a premature stop codon (p.Ser383*). Since *de novo* and ultra-rare psychiatric risk variants generally occur in the heterozygous state [23, 24], we focused on heterozygous mutant mice to model the clinically relevant context. Our mouse model revealed behavioral deficits potentially related to anxiety- and depression-like phenotypes. Analysis of the transcriptome, proteome, and phosphoproteome identified dysregulation in synaptic organization and signaling, neurite morphogenesis, and PKA signaling, which was further corroborated by live-cell imaging of PKA activity and morphometric analysis of mutant neurons.

## Results

### Heterozygous *Akap11* protein-truncating variant reduces *Akap11* mRNA and protein in mice

Using CRISPR-Cas9 genome editing, we introduced a single-nucleotide variant, c.1151C>A; p.Ser384* into exon 7 of mouse *Akap11* transcript (ENSMUST00000123853) (Fig. 1A). Indels in the top five predicted off-target sites were absent in the F2 male founder in PCR gel electrophoresis (Supplementary Fig. 1G) and Sanger sequencing (data not shown). Both heterozygous and homozygous mice were viable. Genotype had a marginal effect on body weight, with the largest difference between heterozygous and homozygous mice (Supplementary Fig. 1H), and significantly affected body length (measured from the nose tip to tail base), with both heterozygous and homozygous mutants shorter than control mice (Supplementary Fig. 1H). RT-qPCR indicated approximately 50% reduction in *Akap11* transcripts in the somatosensory cortex of *Akap11^+/S384*^* mice, using primer pairs targeting regions 5’ or 3’ to the variant (Fig. 1B). Full-length RNA sequencing detected no novel *Akap11* transcript isoforms in the somatosensory cortex of homozygous mutant mice (*Akap11^S384*/S384*^*), as compared to control mice (*Akap11^+/+^*) (Supplementary Fig. 1A). Akap11 protein expression was also significantly reduced in the somatosensory cortex, hippocampus, prefrontal cortex, and striatum of *Akap11^+/S384*^* mice (Fig. 1C-D).

**Fig. 1.**
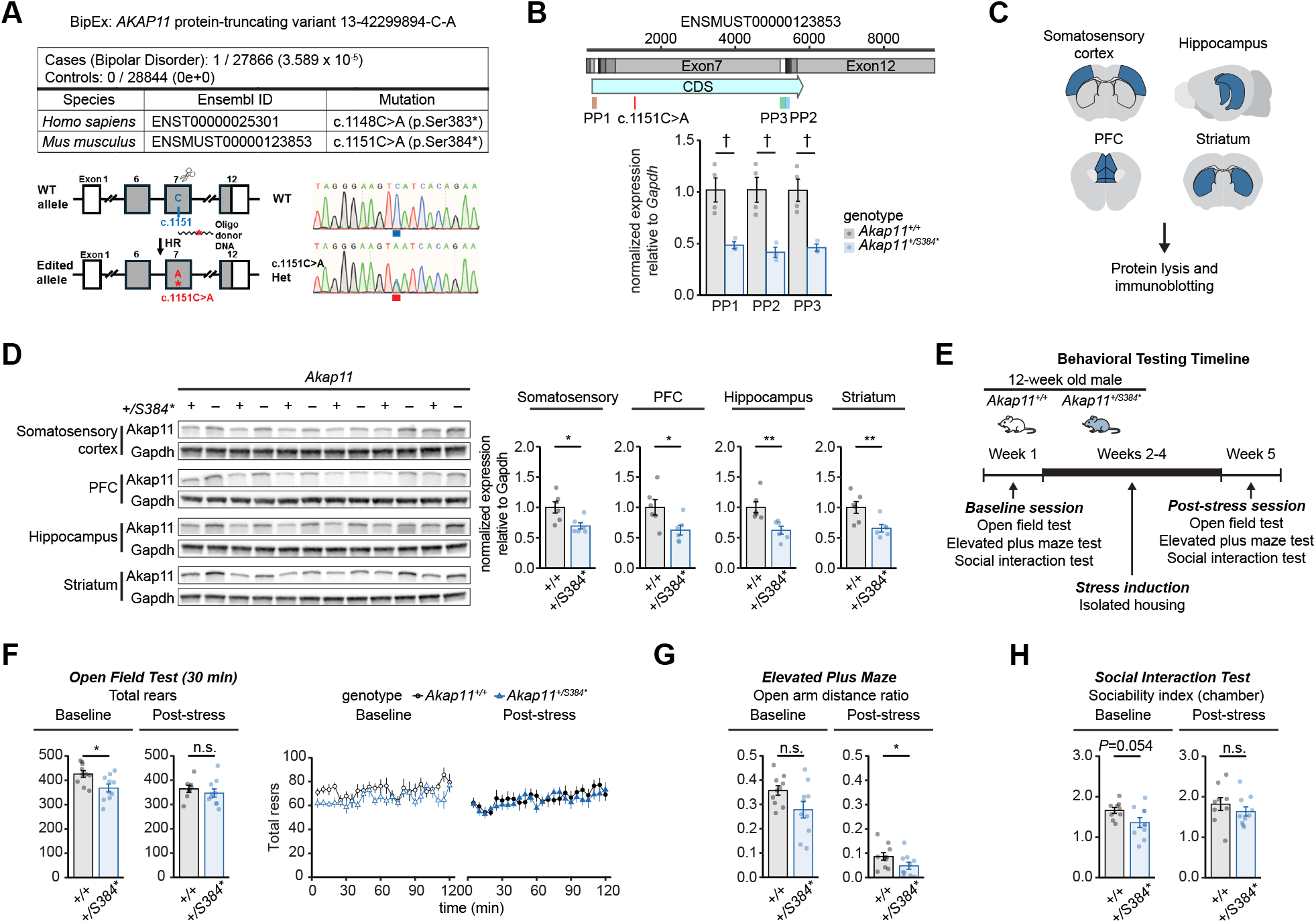
Heterozygous mice carrying a patient-derived ultra-rare Akap11 protein-truncating variant show reduction in Akap11 mRNA and protein, and behavioral deficits related to anxiety- and depression-like phenotypes. A. Schematic of a patient-derived ultra-rare protein-truncating variant (PTV) in *AKAP11* and generation of the knock-in mouse model by CRISPR-Cas9 genome editing. B. Normalized *Akap11* mRNA levels relative to *Gapdh* in *Akap11^+/+^* and *Akap11^+/S384*^* mice. PP1, PP2, and PP3 indicate sequences amplified by three primer pairs. The red bar indicates the position of the single-nucleotide mutation. †*P* = 0.057 via Mann-Whitney U test. Full statistical results are provided in Supplementary Table 1A. C. Schematic showing the brain regions analyzed by immunoblotting for Akap11 protein abundance. Regions analyzed are indicated in blue. D. Immunoblots of Akap11 in four brain regions (left) and quantification (right). Akap11 protein abundance was reduced in the somatosensory cortex (*P* = 0.015), prefrontal cortex (*P* = 0.041), hippocampus (*P* = 0.0022), and striatum (*P* = 0.0087). \**P* ≤ 0.05; \*\**P* ≤ 0.01 via Mann-Whitney U test. Full statistical results are provided in Supplementary Table 1B. E. Timeline of behavioral testing at baseline and after stress induction by isolated housing for three weeks. 10 *Akap11^+/384*^* and 10 *Akap11^+/+^* mice (male, 12 weeks old) completed all behavioral tests. For each measurement, genotype-specific outliers were removed using the 1.5 x interquartile range (IQR) criterion. F. Total rears in open field test during the first 30 min at baseline (bar plots, left) and after three weeks of isolated housing (bar plots, right). At baseline, *Akap11^+/S384*^* mice showed significantly fewer total rears than *Akap11^+/+^* mice (*P* = 0.023). Line plots show the time course of total rears across the 2-h open field test at baseline (line graphs, left) and after isolated housing (line graphs, right). For baseline data, no outliers were removed from the 30-min rearing analysis, and one *Akap11^+/384*^* mouse was removed from the 2-h rearing analysis. For post-stress data, two *Akap11^+/+^* mice were removed as outliers from the 30-min rearing analysis, and no outliers were removed from the 2-h rearing analysis. \**P* ≤ 0.05; n.s. *P* > 0.05 via Mann-Whitney U test. Full statistical results are provided in Supplementary Table 2A. G. Ratio of distance traveled in the open arms of the elevated plus maze at baseline (left) and after isolated housing (right). *Akap11^+/S384*^* mice showed significantly reduced open-arm distance ratio after isolated housing (*P* = 0.043). No outliers were removed from the analysis of open-arm distance ratio. \**P* ≤ 0.05; n.s. *P* > 0.05 via Mann-Whitney U test. Full statistical results are provided in Supplementary Table 2B. H. Chamber-based sociability index in the social interaction test at baseline (left) and after three weeks of isolated housing (right). *Akap11^+/S384*^* mice showed a trend toward reduced sociability (*P* = 0.054) at baseline. One *Akap11^+/+^* mouse was removed as an outlier from the analysis of chamber-based sociability index at baseline and post-stress, respectively. n.s. *P* > 0.05 via Mann-Whitney U test. Full statistical results are provided in Supplementary Table 2C. Histograms show means, and error bars show SEM. *+/+*, *Akap11^+/+^*; *+/S384\**, *Akap11^+/S384*^*.

### Male heterozygous mutant mice exhibit anxiety- and depression-related behavioral deficits

Next, we assessed behavioral consequences of the *Akap11* variant. Since gene-environment interaction is a crucial component in the pathogenesis of neuropsychiatric disorders [25, 26], we evaluated behavioral phenotypes of male *Akap11^+/S384*^* and *Akap11^+/+^* mice at baseline and after isolated housing for three weeks (Fig. 1E), a paradigm to induce mild chronic stress in rodents [27, 28]. In the open field test, *Akap11^+/S384*^* mice showed decreased rearing during the first 30 min at baseline, with unaltered total activity or central ratio (Supplementary Fig. 1B). After three weeks’ isolated housing, the difference in rearing was no longer observed, possibly due to reduced rearing in the *Akap11^+/+^* mice (Fig. 1F). Total activity, central ratio or rears did not differ by genotype in a 2-h open field test (Supplementary Fig. 1B). In the elevated plus maze test, *Akap11^+/S384*^* mice showed a trend of reduced open-arm time ratio at baseline (*P* = 0.052; Supplementary Fig. 1E), and a significantly lower open-arm distance ratio after isolated housing (Fig. 1G), without difference in total distance traveled or zone entries (Supplementary Fig. 1D). Both genotypes showed a preference for closed over the open arms (Supplementary Fig. 1C). In the three-chamber social interaction test, *Akap11^+/S384*^* mice showed a trend of reduced chamber-based sociability index at baseline (*P* = 0.054; Fig. 1H), which was not observed after isolated housing (Fig. 1H). *Akap11^+/S384*^* mice also spent more time in the toy chamber compared to *Akap11^+/+^* mice at baseline, while mouse chamber time and interaction-based sociability index remained unaltered by genotype (Supplementary Fig. 1F). Taken together, heterozygous *Akap11* mutants exhibit behavioral deficits potentially related to anxiety/depression-like phenotypes, with some deficits present at baseline, and others modulated by mild chronic stress.

### Transcriptional changes in the cortex of heterozygous mutant mice highlight alterations in synaptic organization, synaptic signaling, and neuron projection

To identify molecular changes associated with the pathogenic *Akap11* PTV, we performed snRNA-seq of the sensory, medial and lateral neocortical regions from two 12-week-old *Akap11^+/S384*^* mice and three *Akap11^+/+^* littermate controls (Fig. 2A). Clustering of 94693 cells and 21857 genes identified 33 clusters corresponding to layer-specific glutamatergic cell types, major GABAergic cell types, and non-neuronal cell types [29, 30] (Fig. 2B-C; Supplementary Table 3A). Given evidence for substantial transcriptomic, morphological and physiological heterogeneity within cell types in human and mouse brain [30, 31], we further resolved these 33 cell-type clusters into 92 subtypes (Supplementary Table 3A) and evaluated transcriptional changes at multiple resolutions. No significant alterations in cell type or subtype composition between genotypes were detected at false discovery rate (FDR) ≤ 0.05 (Supplementary Fig. 2A-C; Supplementary Table 3B).

**Fig. 2.**
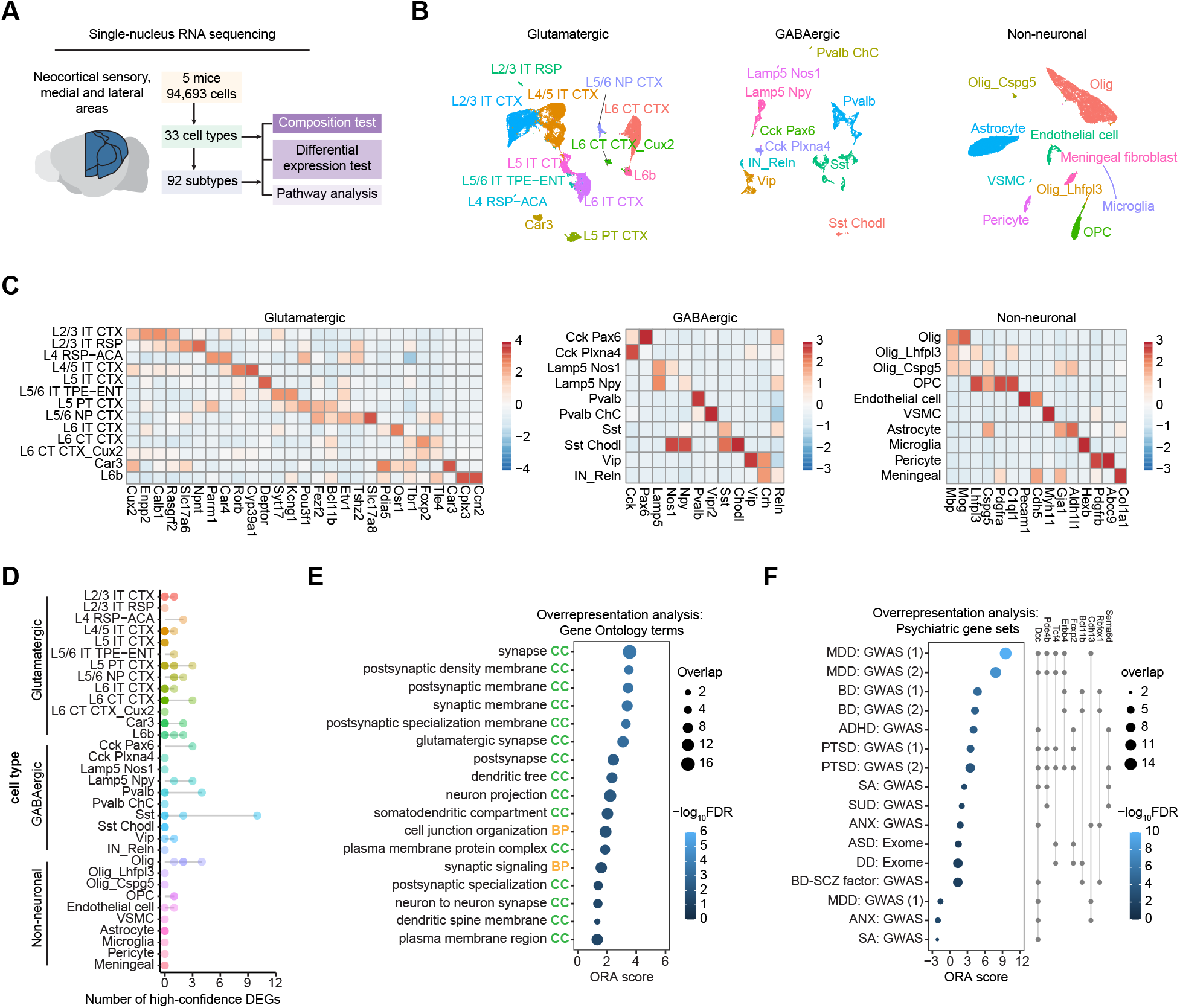
Single-nucleus RNA sequencing revealed transcriptional changes related to synaptic programs and neuron projection in the cortex of heterozygous mutant mice. A. Schematic of single-nucleus RNA sequencing (snRNA-seq) workflow. A total of 33 cell types and 92 subtypes were identified. B. UMAPs of cells annotated into 33 cell types, grouped into glutamatergic, GABAergic, and non-neuronal categories. C. Heatmaps showing mean normalized, log_2_-transformed expression of cell-type markers by the 33 cell types. D. Lollipop plot showing the number of high-confidence differentially expressed genes (DEGs) detected in each of the 92 subtypes, grouped by the cell type. High-confidence DEGs were defined as genes with false discovery rate (FDR) ≤ 0.10 in pseudobulk differential expression analysis using *edgeR* quasi-likelihood negative binomial generalized log-linear model, and nominal *P* ≤ 0.05 in single-cell Poisson-gamma mixed model (PMM), with concordant direction of change between methods. E. Gene ontology (GO) terms overrepresented among upregulated high-confidence DEGs. Significant overrepresentation was defined as FDR ≤ 0.05 via hypergeometric test. The overrepresentation analysis (ORA) score was calculated as −log_10_FDR multiplied by the direction of change, with positive values indicating enrichment among genes upregulated in *Akap11^+/S384*^* mice, and negative values indicating enrichment among downregulated genes. BP, biological process; CC, cellular component. F. Psychiatric disorder risk gene sets overrepresented among extended high-confidence DEGs. Significant overrepresentation is defined as FDR ≤ 0.05 via hypergeometric test. The ORA score is defined as −log_10_FDR multiplied by the direction of change. The matrix grid on the right indicates overlapping genes between each risk gene set and the extended high-confidence DEG list. Only genes present in at least four overrepresented gene sets are shown. Abbreviations: ADHD=attention-deficit hyperactivity disorder, ANX=anxiety disorder, ASD=autism spectrum disorder, BD=bipolar disorder, DD=developmental disorder, MDD=major depressive disorder, PTSD=posttraumatic stress disorder, SA=suicide attempt, SCZ=schizophrenia, and SUD=substance use disorders.

High-confidence differentially expressed genes (DEGs) were determined by FDR ≤ 0.10 in a pseudobulk differential expression (DE) test and nominal *P* ≤ 0.05 in a generalized linear mixed model (GLMM)-based DE test, with concordant direction of change between methods (Supplementary Fig. 2D; SI Methods). These criteria identified 28 DEGs across the 33 cell types (Supplementary Fig. 2E; Supplementary Table 3C), and 61 DEGs across the 92 subtypes (Fig. 2D; Supplementary Fig. 2F; Supplementary Table 3D). Among the 92 subtypes, a somatostatin (SST)-expressing subtype contained the largest number of DEGs, all upregulated in the *Akap11^+/S384*^* mice (Fig. 2D; Supplementary Fig. 2F). This subtype showed elevated *Hpse* expression, suggesting potential correspondence to non-Martinotti SST interneurons in cortical layer 4 (L4) and L5a [32].

Overrepresentation analysis using Gene Ontology (GO) terms and synapse-specific GO (SynGO) pathways was performed for the high-confidence DEGs across the 92 subtypes (Fig. 2E; Supplementary Fig. 2G-H; Supplementary Tables 3E-F). Upregulated DEGs were significantly enriched for GO pathways related to synaptic organization and signaling (FDR ≤ 0.05; Fig. 2E). Consistent with these alterations, the upregulated DEGs were also enriched for a subset of SynGO pathways, predominantly associated with the postsynaptic compartment or general synapse categories (Supplementary Fig. 2G-H). In addition, upregulated DEGs were enriched for GO terms related to neurite morphogenesis, including “dendritic tree”, “neuron projection”, “somatodendritic compartment”, and “dendritic spine membrane” (Fig. 2E). Downregulated DEGs did not show significant enrichment for GO or SynGO terms.

We next evaluated whether transcriptional changes in *Akap11^+/S384*^* mice were associated with risk genes for neuropsychiatric disorders based on GWAS or WES studies (Supplementary Table 3G). Using an extended DEG set defined by nominal significance in the pseudobulk DE test (*P* ≤ 5 x 10^-4^) and the GLMM-based test (*P* ≤ 0.05) with concordant direction of change, we identified an enrichment for risk genes associated with mood and psychotic disorders, including major depressive disorder, bipolar disorder, and schizophrenia, among other disorders (Fig. 2F; Supplementary Table 3H). Several DEGs were implicated across multiple disorders, including *Dcc*, which encodes the Netrin-1 receptor that regulates axon guidance [33], and has cross-disorder effects on bipolar disorder and schizophrenia [5] (Fig. 2F). The extended DEGs were also enriched for curated DisGeNET gene sets related to mood, psychotic, and developmental disorders [34] (Supplementary Fig. 2I; Supplementary Table 3I).

### Shared and cell type-specific transcriptional changes in the cortex of heterozygous mutant mice

We next analyzed potential shared and cell type-specific transcriptional changes. *Spon1*, which encodes the extracellular matrix adhesion molecule F-spondin that regulates neurite outgrowth and cell migration [35, 36], was downregulated in a parvalbumin-expressing interneuron subtype (Fig. 3A). We also observed dysregulation of four cell adhesion molecule genes (*Alcam*, *Cdh12*, *Nxph1* and *Cdh13*) across four deep-layer projection neuron subtypes that innervate distinct cortical and subcortical targets (Fig. 3A). Given their roles in neurite outgrowth and synapse formation [37–39], these changes suggest altered neuron projection and connectivity in the heterozygous cortex. In addition, *Sgk1* was elevated in two mature oligodendrocyte subtypes (Fig. 3A; Supplementary Table 3D), and at the oligodendrocyte cell-type level (Supplementary Table 3C). *Sgk1* encodes the serum and glucocorticoid-regulated kinase 1, and exposure to stress increases *Sgk1* expression in oligodendrocytes and alters oligodendrocyte morphology in mice [40, 41].

**Fig. 3.**
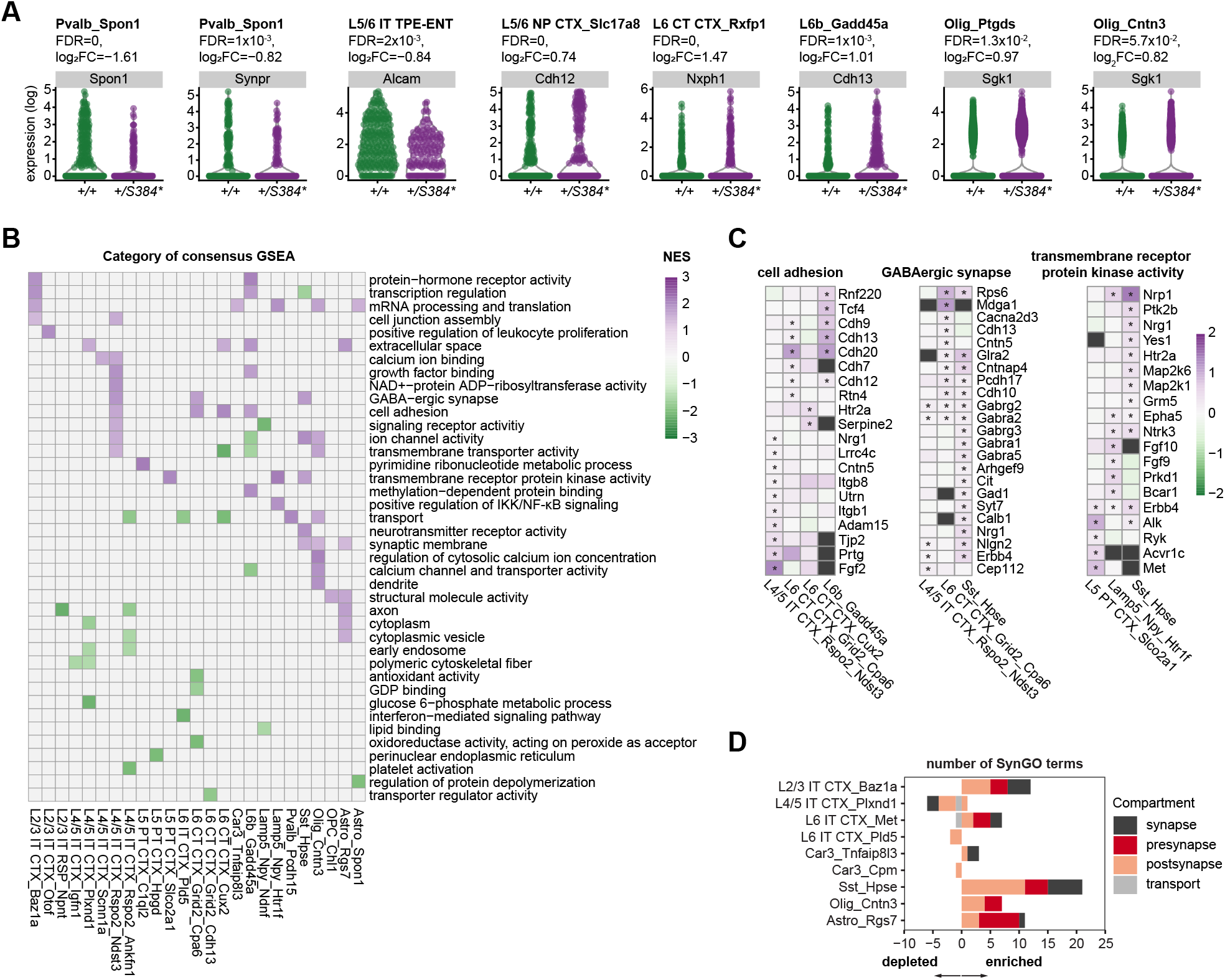
snRNA-seq revealed shared and cell type-specific transcriptional changes in the cortex of heterozygous mutant mice. _A._ Violin plots showing normalized, log_2_-transformed expression of select differentially expressed genes (DEGs) between *Akap11^+/+^* and *Akap11^+/S384*^* mice. *+/+*, *Akap11^+/+^*; *+/S384\**, *_Akap11+/S384*_*. B. Heatmap showing mean normalized enrichment score (NES) for GO term categories across subtypes with at least 5,000 genes meeting the minimum expression criteria. Positive and negative NES values indicate enrichment and depletion in *Akap11^+/S384*^* mice, respectively. Significance was defined as FDR ≤ 0.05 via permutation test. C. Heatmaps showing ranks of leading-edge genes for selected GO categories. Gene ranks were calculated as −log_10_*P* x sign(log_2_ fold change) from the pseudobulk differential expression test. Positive and negative ranks indicate up- and down-regulation, respectively, in the *Akap11^+/S384*^* mice. Asterisks indicate leading-edge genes for the corresponding subtype and pathway category. Only genes with *P* ≤ 0.05 in the pseudobulk differential expression test in at least one relevant subtype are shown. D. Number of significantly enriched or depleted SynGO terms in each subtype. Significance was defined as FDR ≤ 0.05 via permutation test.

To dissect the biological pathways affected by the *Akap11* PTV, we performed gene set enrichment analysis (GSEA) for each subtype. Genes were ranked by −log_10_*P* x sign(log_2_ fold change). To ensure validity of the permutation test, GSEA was performed only for clusters with at least 5000 genes passing expression-level filtering. We observed shared and cell type-specific enrichment or depletion of 124 GO pathways across subtypes, which were summarized into 40 categories based on semantic similarity and ancestor relationships (Fig. 3B; Supplementary Table 4A; SI Methods). GO terms related to cell adhesion were enriched in *Akap11^+/S384*^* mice across one L4/5 intratelencephalic (IT) subtype, two L6 corticothalamic (CT) subtypes, and one L6b subtype (Fig. 3B). Genes driving enrichment included the classic and atypical cadherins (Fig. 3C). We also observed enrichment of GO terms related to GABAergic signaling in L4/5 IT, L6 CT, and SST neuron subtypes, as well as enrichment of pathways related to transmembrane receptor protein kinase activity in L5 pyramidal tract (PT), Lamp5, and SST neuron subtypes (Fig. 3B-C). GSEA using synapse-specific SynGO pathways revealed that the mutant was predominantly associated with upregulation of synaptic gene sets (Fig. 3D; Supplementary Table 4B). Some synapse-related terms were also enriched in non-neuronal subtypes, including oligodendrocyte and astrocyte subtypes (Fig. 3B, D). Similar observations have been made in one prior transcriptomic study of these cell types [42], which may reflect non-specific annotation of some GO terms.

### Patient-derived *Akap11* PTV alters the proteome in the cortex of heterozygous mutant mice

We used TMT-based LC-MS/MS to profile the total proteome and phosphoproteome of the neocortex of five *Akap11^+/S384*^* and four *Akap11^+/+^* male mice at 12 weeks of age (Fig. 4A). Database search of the total proteome identified 8742 protein isoforms. Due to the small sample size, we used a nominal *P* value threshold (*P* **≤** 5 x 10^-4^) in combination with effect size filtering (absolute log_2_ fold change ≥ 0.10) to prioritize differentially abundant proteins (DAPs; Fig. 4B; Supplementary Table 5A). Nine DAPs showed increased abundance in *Akap11^+/S384*^* mice, including Prkar1a and Prkar1b (Fig. 4B), the type I PKA regulatory subunits RIα and RIβ, respectively. Prkaca and Prkacb, the PKA catalytic subunits α (Cα) and β (Cβ), were also increased in *Akap11^+/S384*^* mice (Fig. 4B). In contrast, the type II PKA regulatory subunits RIIα and RIIβ were not differentially abundant between genotypes (Supplementary Table 5A). In immunoblotting, PKA RIα and RIβ protein levels were increased by 69.8% and 36.3%, respectively, and the catalytic subunits Cα and Cβ were increased by 14.5% and 24.3%, respectively, with large effect sizes (*r* ≥ 0.79; Fig. 4D). Type II regulatory subunits RIIα and RIIβ remained unaltered in the cortex of *Akap11^+/S384*^* mice (Fig. 4D). Our results thus suggested selective elevation of type I PKA regulatory subunits and PKA catalytic subunits in the cortex of *Akap11^+/S384*^* mice.

**Fig. 4.**
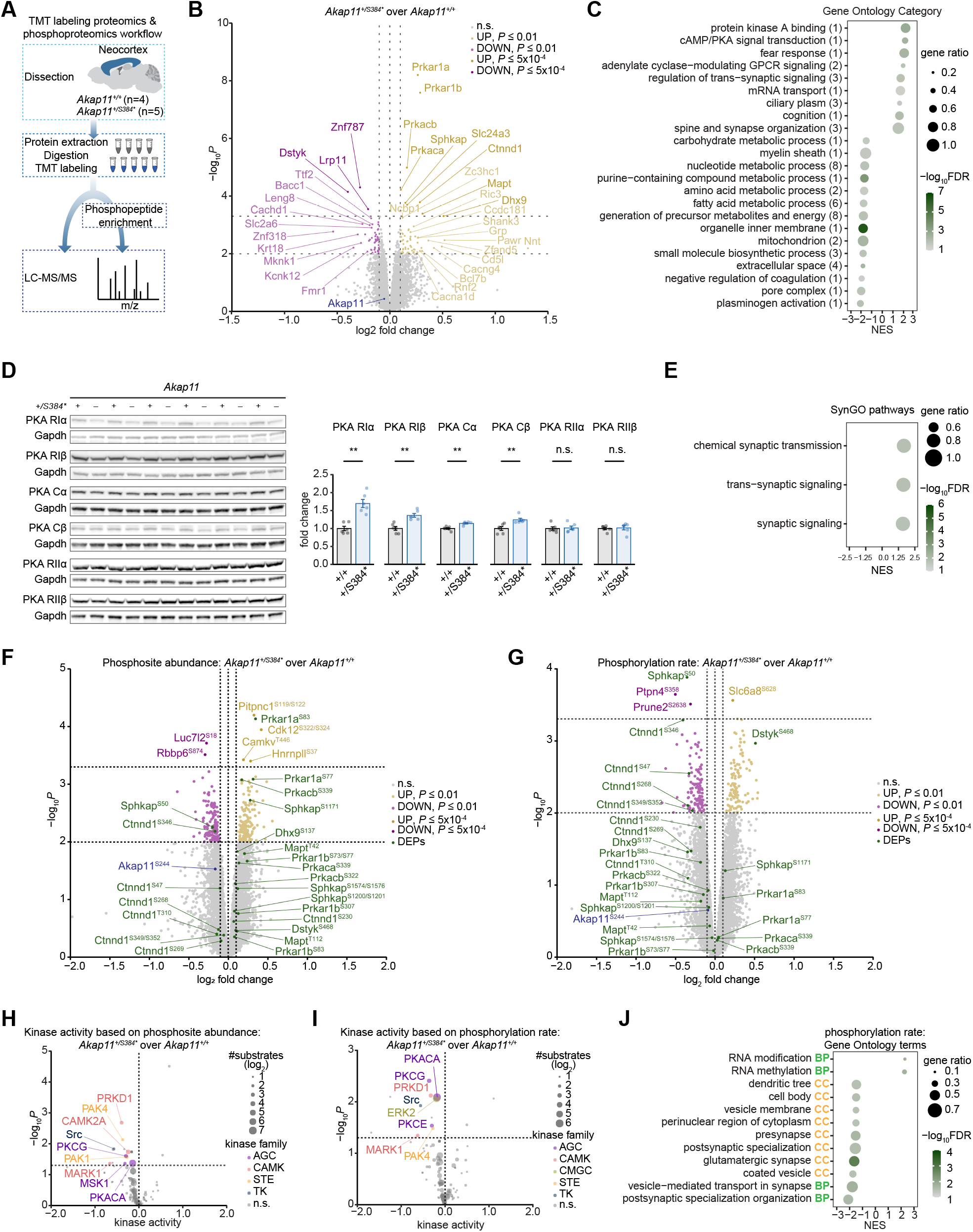
The patient-derived *Akap11* PTV alters the cortical proteome and phosphoproteome in heterozygous mutant mice. A. Overview of the TMT labeling proteomics and phosphoproteomics workflow using cortical tissues from five *Akap11^+/S384*^* and four *Akap11^+/+^* mice. B. Volcano plot showing the differentially abundant proteins (DAPs) and Akap11 (dark blue). *P* values were estimated via moderated *t*-tests using *limma*. Positive and negative log_2_ fold changes indicate increased and decreased protein abundance, respectively, in *Akap11^+/S384*^* mice. Proteins meeting nominal *P* ≤ 5 x 10^-4^ are shown in gold for increased abundance (nine proteins), and purple for decreased abundance (three proteins). Proteins meeting nominal *P* ≤ 0.01 are shown in light gold or light purple. Horizontal dashed lines indicate nominal *P* cutoffs of 5 x 10^-4^ (upper) and 0.01 (lower). Vertical dashes lines indicate log_2_ fold changes of -0.1, 0.0 and 0.1. C. Dot plot showing GO term categories significantly enriched or depleted in *Akap11^+/S384*^* mice via permutation test. Numbers in parentheses indicate the number of GO terms assigned to each category. Positive and negative normalized enrichment scores (NES) indicate enrichment and depletion, respectively, in *Akap11^+/S384*^* mice. Category-level NES values represent the mean NES of GO terms within each category. Gene ratio was computed as the proportion of pathway genes represented in the ranked protein list. Significance was defined as FDR ≤ 0.05. D. Immunoblots and quantification of type I PKA regulatory subunits RIα and RIβ, PKA catalytic subunits Cα and Cβ, and type II PKA regulatory subunits RIIα and RIIβ. Difference between genotypes were significant for RIα (*P* = 0.0022), RIβ (*P* = 0.0022), Cα (*P* = 0.0043), and Cβ (*P* = 0.0022), but not for RIIα (*P* = 0.70), or RIIβ (*P* = 0.31). \*\**P* ≤ 0.01, *n.s.*, *P* > 0.05 via Mann-Whitney U test. *+/+*, *Akap11^+/+^*; *+/S384\**, *Akap11^+/S384*^*. Full statistical results are shown in Supplementary Table 5B. E. Gene set enrichment analysis (GSEA) of SynGO pathways via permutation test. Positive and negative NES values indicate enrichment and depletion, respectively, in *Akap11^+/S384*^* mice. Gene ratio was computed as the proportion of pathway genes represented in the ranked protein list. Significance was defined as FDR ≤ 0.05. F-G. Volcano plots showing the genotype-dependent differences in phosphosite abundance (F), and phosphorylation rate (G), defined as phosphosite abundance normalized to the abundance of the corresponding master protein. Differential expression was defined as nominal *P* ≤ 5 x 10^- 4^, and absolute log_2_ fold change ≥ 0.10 in moderated *t-*tests via *limma*. Positive and negative log_2_ fold changes indicate increased and decreased phosphosite abundance or phosphorylation rate, respectively, in *Akap11^+/S384*^* mice-. Gold and purple indicate phosphosites increased or decreased at nominal *P* ≤ 5 x 10^-4^, respectively light gold and light purple indicate phosphosites increased or decreased at nominal *P* ≤ 0.01, respectively. Phosphosites associated with differentially abundant proteins in Fig. 4B are shown in green, and Akap11 phosphosites are shown in blue. Horizontal dashed lines indicate nominal *P* thresholds at 5 x 10^-4^ (upper) or 0.01 (lower). H-I. Volcano plots showing predicted kinase activity based on phosphosite abundance (H) or phosphorylation rate (I). Kinase activity scores and *P* values were estimated by the *RoKAI* method, applied to −log_10_*P* x sign(log_2_ fold change). Positive and negative kinase activity scores indicate increased and decreased kinase activity, respectively, in *Akap11^+/S384*^* mice. Kinases were considered significant was at *P* ≤ 0.05 with at least three substrates present. J. GSEA of GO terms applied to phosphosites ranked by their differential phosphorylation-rate scores, calculated as −log_10_*P* x sign(log_2_ log fold change). Positive and negative NES scores indicate pathways associated with proteins harboring increased or decreased phosphorylation rate, respectively, in *Akap11^+/S384*^* mice. *P* values were estimated via permutation test, and significance was defined as FDR ≤ 0.05. Gene ratio was computed as the percentage of pathway genes present in the phosphosite dataset.

We also observed an increase in Ctnnd1 (p120-catenin), encoded by a risk gene for multiple psychiatric disorders, including major depressive disorder and schizophrenia [43]; Sphkap, an AKAP that preferentially binds to PKA RIα [44]; and Mapt, the tau protein which is linked to neurodegenerative diseases [45], among other proteins (Fig. 4B). Three DAPs showed reduced abundance in *Akap11^+/S384*^* mice. With a less stringent criterion (*P* ≤ 0.01 and absolute log_2_ fold change ≥ 0.10), 117 DAPs were detected in the *Akap11^+/S384*^* neocortex, including 62 increased and 55 decreased proteins (Fig. 4B).

GSEA of the total proteome revealed 56 significantly enriched or depleted GO terms, summarized into 23 categories (Fig. 4C, Supplementary Table 5C). Terms related to “protein kinase A binding”, “cAMP/PKA signal transduction”, and “adenylate cyclase-modulating GPCR signaling” were enriched in *Akap11^+/S384*^* mice (Fig. 4C), suggesting altered PKA signaling.

Consistently, Reactome pathways related to GPCR signaling were enriched in *Akap11^+/S384*^* mice (Supplementary Fig. 3A; Supplementary Table 5D). The enrichment of PKA signaling and adenylate cyclase-modulating GPCR signaling pathways was primarily driven by increased abundance of PKA holoenzyme components, adenylate cyclase-associated proteins, and PKA-interacting proteins (Supplementary Fig. 3B). Synapse-related pathways were also enriched (Fig. 4C, E; Supplementary Tables 5C, E), consistent with the transcriptomic findings (Fig. 2F). In contrast, pathways associated with myeline sheath, biomolecule metabolism, and mitochondrial processes were depleted in *Akap11^+/S384*^* mice (Fig. 4C; Supplementary Fig. 3A-B).

### Altered phosphoproteome and kinase activity in the cortex of heterozygous mutant mice

The patient-derived *Akap11* PTV selectively increased type I PKA regulatory subunits RIα/β and PKA catalytic subunits Cα/β. Since a change in PKA isoform abundance may affect downstream PKA-dependent phosphorylation, we asked how the overall phosphoproteomic landscape and PKA kinase activity would be altered.

10709 phosphosites were identified from the phospho-enriched fractions of the LC-MS/MS data. Nine were differentially abundant (seven increased, and two decreased) in *Akap11^+/S384*^* neocortex (*P* ≤ 5 x 10^-4^ and absolute log_2_ fold change ≥ 0.10; Fig. 4F, Supplementary Table 6A). With a less stringent cutoff (*P* ≤ 0.01 and absolute log_2_ fold change ≥ 0.10), 164 increased and 161 decreased phosphosites were identified (Fig. 4F). For the 12 DAPs in the bulk proteome (Fig. 4B), we detected an increase in Prkar1a^S83^ at *P* ≤ 5 x 10^-4^, Prkar1a^S77^, Prkacb^S339^, and Sphkap^S1171^ at *P* ≤ 0.01, and a decrease in Sphkap^S50^ and Ctnnd1^S346^ at *P* ≤ 0.01 (Fig. 4F).

Phosphorylation of PKA RIα at the two conserved sites S83 and S77 increases binding of RIα to Cα and inhibits PKA activity [46]. We thus hypothesized that PKA activity would be reduced in *Akap11^+/S384*^* mice and estimated kinase activity from the phosphorylation state of substrates using kinase-substrate annotations in the PhosphoSitePlus [47] and the Signor databases [48]. Using a *Z* test-based inference method [49], we identified significant decrease in PKA (PKACA) kinase activity in *Akap11^+/S384*^* neocortex with 66 overlapping substrates (Fig. 4H; Supplementary Fig. 3D; Supplementary Tables 6B-C). The trend of reduced PKA kinase activity was confirmed using an alternative kinase-substrate database and inference test (*P* = 0.057; Supplementary Fig. 3C; Supplementary Table 6D; SI Methods). GSEA revealed that the myelin sheath pathway was associated with proteins containing phosphosites with reduced abundance (Supplementary Fig. 3G; Supplementary Table 6E).

Phosphorylation rate was assessed by normalizing phosphosite abundance to the corresponding bulk proteome abundance. Four out of 9565 phosphosites showed altered phosphorylation rate at *P* ≤ 5 x 10^-4^ and absolute log_2_ fold change ≥ 0.10 (Fig. 4G; Supplementary Table 6F), while 263 were altered at the less stringent cutoff (*P* ≤ 0.01 and absolute log_2_ fold change ≥ 0.10) (Fig. 4G). Among the 12 DAPs, Sphkap showed reduced phosphorylation rate at S50 (Fig. 4G) despite increased total protein abundance (Fig. 4B); Ctnnd1 showed reduced phosphorylation at five sites, and Dstyk showed increased phosphorylation at one site (Fig. 4G). Kinase-substrate analysis based on phosphorylation rate indicated reduced PKA (PKACA) activity (Fig. 4I; Supplementary Fig. 3E-F; Supplementary Tables 6G-I).

GSEA showed that proteins associated with reduced phosphorylation rates in *Akap11^+/S384*^* mice were enriched for synapse-related terms (Fig. 4J; Supplementary Table 6J). SynGO analysis confirmed enrichment of synaptic pathways among phosphosites with reduced phosphorylation rates (Supplementary Fig. 3H; Supplementary Table 6K). Ctnnd1 and Prune2, both containing phosphosites with reduced phosphorylation rates in *Akap11^+/S384*^* mice (Fig. 4G), were among leading-edge contributors to synapse-related pathways (Supplementary Table 6K). The GO pathway “dendritic tree” was also associated with proteins harboring phosphosites with reduced phosphorylation rates (Fig. 4J). Thus, synapse- and neuron projection-related processes appeared as recurrent dysregulated themes across the transcriptome, proteome, and phosphoproteome of the neocortex of *Akap11^+/S384*^* mice. In addition, vesicle-related pathways were enriched among proteins containing phosphosites with reduced phosphorylation rates (Fig. 4J).

### Attenuated stimulus-dependent PKA activity in excitatory neurons of heterozygous mutant mice

Our proteomic analysis indicated selective elevation of type I, but not type II, PKA regulatory subunits, together with increased PKA catalytic subunits and reduced phosphorylation of predicted PKA substrates in the neocortex of heterozygous mutant mice (Fig. 4), suggesting altered PKA signaling. Cortical neurons derived from E16.5 *Akap11^+/S384*^* embryos showed increased PKA RIα immunoreactivity at DIV 21-22 (Fig. 5A-B). To directly assess PKA kinase activity, we expressed the excitation-ratiometric PKA sensor ExRai-AKAR2 [50] under the Camk2a promoter in cultured cortical neurons from E16.5 *Akap11^+/S384*^* and *Akap11^+/+^* embryos (Fig. 5C-D). Sensor specificity and bidirectional sensitivity was confirmed in *Akap11^+/+^* neurons using forskolin and rolipram, and/or the PKA inhibitor H-89 (Supplementary Fig. 4A-B).

**Fig. 5.**
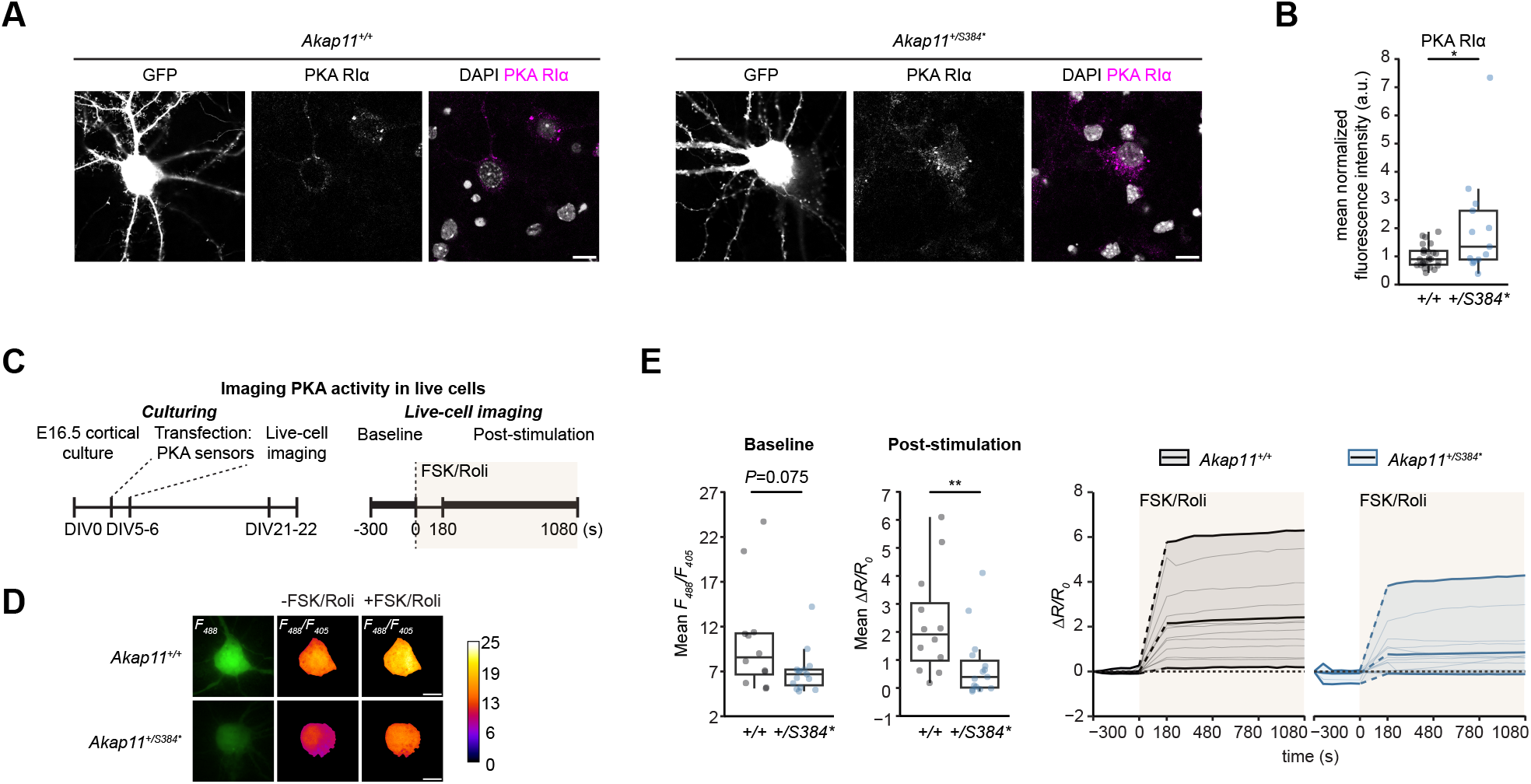
Live-cell imaging reveals attenuated stimulus-dependent PKA activity in excitatory cortical neurons from heterozygous mutant mice. A. Representative images of cortical neurons transfected with pCAG-GFP at days *in vitro* (DIV) 4-6 and immunostained for PKA RIα at DIV21-22. Scale bars, 10 µm. B. Quantification of PKA RIα immunofluorescence in the soma of *Akap11^+/S384*^* and *Akap11^+/+^* neurons. *Akap11^+/S384*^* neurons showed elevated PKA RIα immunofluorescence compared to *Akap11^+/+^* neurons (*Akap11^+/+^*: N = 5 embryos, n = 26 neurons; *Akap11^+/S384*^*: N = 6 embryos, n = 13 neurons; *P* = 0.047). \**P* ≤ 0.05 via Mann-Whitney U test. Full statistical results are shown in Supplementary Table 7A. C. Experimental timeline for cortical neuron culturing, transfection with the PKA sensor, and live- cell imaging. D. Representative PKA sensor signals in an *Akap11^+/+^* neuron (upper panels) and an *Akap11^+/S384*^* neuron (lower panels) at DIV22. For each neuron, the *F_488_* signal after forskolin/rolipram stimulation (left), the *F_488_/F_405_* ratio at baseline (-FSK/Rol; middle), and the *F_488_/F_405_* ratio after stimulation (+FSK/Roli; right) were shown. Images are shown from the first time point of each condition. Scale bar, 10 µm. FSK, forskolin. Roli, rolipram. E. Quantification of PKA activity. Boxplots show the mean baseline *F_488_/F_405_* ratio (left) and the mean normalized change in *F_488_*/*F_405_* ratio (mean *ΔR/R_0_*) in response to forskolin/rolipram stimulation (right), where *R* = *F_488_*/*F_405_*, *R_0_* is the ratio at the first baseline time point, and *ΔR* is the change in *R* relative to *R_0_*. Baseline PKA activity showed a trend toward reduction in *Akap11^+/S384*^* neurons compared to *Akap11^+/+^* neurons (*P* = 0.075), whereas stimulus-evoked PKA activity was significantly attenuated in *Akap11^+/S384*^* neurons (*Akap11^+/+^*: N = 2 embryos, n = 12 neurons; *Akap11^+/S384*^*: N = 3 embryos, n = 15 neurons; *P* = 0.0074). \*\**P* ≤ 0.01 via Mann- Whitney U test. For boxplots, the bottom edge, midline, and top edge represent the first quantile, median, and third quantile, respectively. Whiskers represent 1.5 x interquartile range from the box edges. Line plots show the time course of *ΔR/R_0_* for *Akap11^+/+^* neurons (left) and *Akap11^+/S384*^* neurons (right). Upper, middle, and lower thick solid lines indicate maximum, mean, and minimum *ΔR/R_0_*, respectively. Thin solid lines indicate traces of individual neurons. Dashed lines indicate a 3-min interval during drug application, when imaging was paused.Horizontal dotted lines indicate *ΔR/R_0_* = 0. Full statistical results are shown in Supplementary Table 7C.

In DIV22 cortical excitatory neurons, there was a marginal reduction in baseline PKA activity, indicated by the excitation ratio *F_488_*/*F_405_*, in *Akap11^+/S384*^* neurons compared to *Akap11^+/+^*neurons (*P* = 0.075; Fig. 5E). Stimulus-evoked PKA activity was quantified as *ΔR/R_0_*, where *R* = *F_488_*/*F_405_*, *R_0_* is the ratio at the first baseline time point, and *ΔR* is the change in *R* relative to *R_0_*. Stimulation with forskolin (25 µM) and rolipram (2 µM) increased PKA activity in *Akap11^+/+^* neurons by approximately 2.30 folds, whereas the response was attenuated in *Akap11^+/S384*^* neurons (increased by 0.80 folds) (Fig. 5E). Thus, stimulus-dependent PKA activity was functionally impaired in cortical excitatory neurons from *Akap11^+/S384*^* mice.

### Impaired nascent neurite development in cortical neurons of heterozygous mutant mice

Neuron projection and neurite morphogenesis emerged recurrently as dysregulated pathways in the cortical transcriptome, proteome and phosphoproteome of *Akap11^+/S384*^* mice, in addition to attenuated stimulus-dependent PKA signaling (Fig. 2-5). Given the role of PKA in growth cone guidance and dendrite outgrowth [51, 52], we hypothesized that the patient-derived *Akap11* PTV disrupts neurite development in cortical neurons.

To test this hypothesis, we analyzed DIV6 neurite morphology of cortical neurons from E16.5 *Akap11^+/+^* and *Akap11^+/S384*^* embryos. Nascent axons were identified as Tau-1-positive/MAP2-positive processes, and dendrites as Tau-1-negative/MAP2-positive processes. Soma area, dendritic tree number, and primary axon length did not differ by genotype (Fig. 6A-B). In contrast, *Akap11^+/S384*^* neurons showed significantly reduced average dendritic length (*Akap11^+/+^* neurons: 55.68 µm, *Akap11^+/S384*^* neurons: 40.05 µm; Fig. 6A-B), suggesting impaired early dendritic growth.

**Fig. 6.**
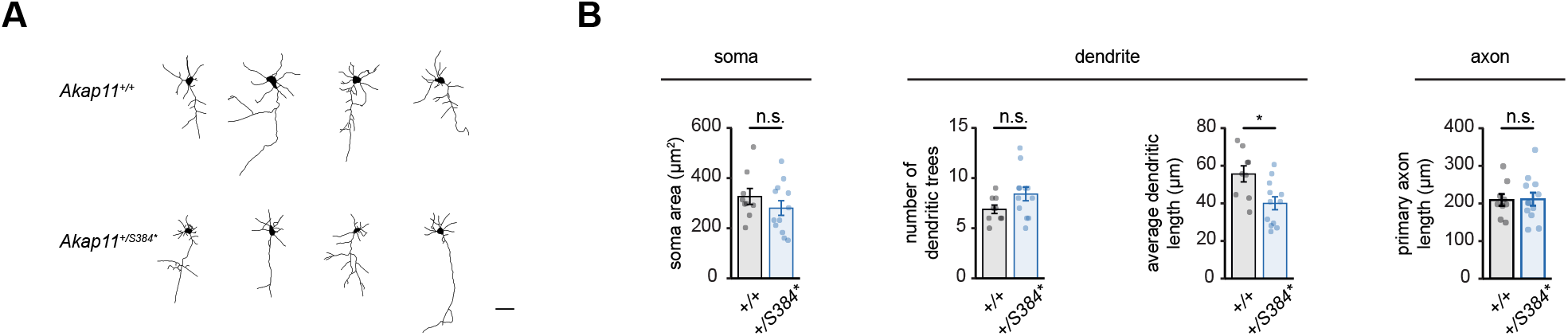
The patient-derived *Akap11* PTV disrupts nascent dendrite morphogenesis. A. Tracings of representative DIV6 cortical neurons in *Akap11^+/+^* and *Akap11^+/S384*^* cultures. Scale bar, 50 µm. B. Quantitative analysis of neuronal morphology at DIV6. *Akap11^+/S384*^* neurons showed no significant differences in soma area (*P* = 0.38), number of dendritic trees (*P* = 0.11), and length of primary axon (*P* = 1.00), but exhibited significantly reduced average dendritic length (*P* = 0.018). *Akap11^+/+^*: N = 2 embryos, n = 12 neurons; *Akap11^+/S384*^*: N = 2 embryos, n = 9 neurons. \**P* ≤ 0.05, n.s., *P* > 0.05 via Mann-Whitney U test. Full statistical results are shown in Supplementary Table 8A.

## Discussion

Recent WES studies have identified ultra-rare PTVs associated with bipolar disorder and schizophrenia [10, 11], but their pathogenic mechanisms remain unclear. Here we focus on an ultra-rare PTV in *AKAP11* identified in a patient with bipolar disorder, and engineered a mouse model with a homologous variant. In heterozygous mutant mice, the variant reduced *Akap11* mRNA and protein levels, whereas the homozygous mutant mice showed no detectable stable novel *Akap11* mRNA isoforms. The current study extends findings from prior *Akap11* KO models in several important ways. First, by modeling a clinically observed single-nucleotide PTV, we showed that one mutant allele was sufficient to produce behavioral abnormalities and PKA dysregulation consistent with prior exon-deletion *Akap11* loss-of-function models. Second, proteomic and transcriptomic analyses identified convergent changes in synapse- and neuron projection-related pathways. Third, functional analyses demonstrated attenuated stimulus-dependent PKA signaling and impaired dendritic morphogenesis in cortical neurons, highlighting potential mechanistic pathways that may be therapeutically relevant.

Heterozygous mutant mice showed reduced rearing in the open field test and impaired sociability at baseline. Isolated housing for three weeks further reduced open-arm activity in the elevated plus maze (Fig. 1). Abnormalities in these measures are linked to anxiety- and depression-like phenotypes in rodents [53–56]. Our results align with prior heterozygous *Akap11* KO models, which showed reduced open-field rearing, sucrose preference, elevated plus-maze open-arm activity, and light-box entries, as well as increased tail-suspension immobility [20, 21]. However, one limitation is that neither our model nor prior KO models (either homozygous or heterozygous) exhibited clear mania-like behaviors, such as hyperactivity and reduced anxiety [57, 58]. Thus, *Akap11* loss-of-function may be sufficient to produce anxiety/depression-related abnormalities but insufficient to capture the full behavioral spectrum of bipolar disorder [59, 60], which likely requires additional polygenic and environmental contributors.

Discrepancies exist regarding Akap11 and PKA subunit expression between our findings and a prior study using an *Akap11* KO model. We detected reduced Akap11 protein expression in four cortical and subcortical regions at 12 weeks of age (Fig. 1D), whereas Song et al. [20], using *Akap11* exon 6-7 deletion mice, showed Akap11 reduction homozygous KO cortical lysates but less consistent changes in the heterozygous KO. We also observed selective elevation in type I PKA regulatory subunits RIα and RIβ, and PKA catalytic subunits Cα and Cβ, consistent with changes in the homozygous KO in Song et al. [20], but differing from their heterozygous KO results, where the elevation in RIα and Cα was not robust. Possible technical explanations are the use of different statistical tests, and the use of β-actin as a loading control in Song et al. [20].

Since AKAP11 and PKA interact with and regulate the actin cytoskeleton [61], normalization to β-actin could potentially mask changes in Akap11 and PKA subunit abundance. On the other hand, extending the synaptic proteomic findings in Song et al. [20], which showed increased type I PKA regulatory subunits and PKA catalytic subunits in the heterozygous KO, our data further showed elevation of these subunits in cortical total lysates from heterozygous mutant mice carrying the patient-derived *Akap11* PTV. As such, we demonstrated that the patient-derived *Akap11* PTV in the heterozygous state is sufficient to reduce Akap11 protein abundance and alter the expression of specific PKA subunits.

Prior *in vitro* PKA substrate phosphorylation assays consistently reported increased basal PKA activity in synaptic fractions of the homozygous KO, whereas changes the cytosolic fractions or total lysates, particularly in the heterozygous KO, remained less consistent [19, 20]. These assays primarily measured constitutive activity, leaving stimulus-dependent PKA dynamics in a cellular context largely unknown. Using time-lapse PKA sensor imaging in the soma of cultured cortical excitatory neurons, we demonstrated a trend of reduced basal PKA activity and attenuated forskolin/rolipram-evoked PKA activation in heterozygous mutant neurons. Stimulus-dependent PKA signaling regulates neural plasticity by modulating phosphorylation, trafficking and activation of synaptic proteins, as well as neurotransmitter release, and cAMP response element-binding protein (CREB)-dependent gene expression [62, 63]. Thus, impaired stimulus-dependent PKA dynamics may undermine synaptic plasticity, affecting long-term potentiation (LTP), learning, and memory. Consistent with this hypothesis, impaired LTP-like cortical plasticity has been implicated in schizophrenia and mood disorders [64, 65].

In Song et al. [20], *in vivo* FLIM-based PKA sensor recordings showed elevated basal PKA activity, and enhanced response magnitude to D1 or D2 receptor antagonists in the striatum of homozygous KO mice, but no changes in the heterozygous KO mice. The inconsistency with our findings may reflect cell type-specific PKA dynamics in the striatum and cortex [66, 67].

Global administration of D1/D2 receptor antagonist may also engage diverse cell types and circuits due to broad distribution of D1/D2 receptors [68]. In contrast, by expressing a PKA sensor in cultured cortical excitatory neurons and bath applying forskolin/rolipram, our approach provided a cell type-specific, direct readout of PKA dynamics. A limitation of our study is that PKA sensor imaging focusing on the soma did not inform PKA dynamics at synapses or in neuronal processes. Current PKA sensors [50, 69] also cannot distinguish isoform-specific PKA signaling, which may require direct assays of PKA RIα-Cα dissociation dynamics in future studies.

## Conclusions

In summary, a heterozygous patient-derived ultra-rare PTV in *Akap11* produces behavioral phenotypes reminiscent of key aspects of psychiatric disorders, and recapitulates molecular changes observed in prior loss-of-function models, with additional evidence for dysregulated stimulus-dependent PKA signaling and impaired morphogenesis in cortical neurons. These data support *Akap11* haploinsufficiency as a potential pathogenic mechanism. Given that *de novo* and ultra-rare coding variants in psychiatric disorders are presumed to act in the heterozygous state [23, 24], this genetically precise model provides a clinically relevant system for investigating how rare coding variants may underlie psychiatric disorders.

## Materials and Methods

### Animals

#### Generation of knock-in mouse line

Heterozygous *Akap11^+/S384*^* mice were generated using a CRISPR/Cas9-mediated homologous recombination strategy on the C57BL6/J genetic background (The Jackson Laboratory, Bar Harbor, ME, USA). The design of the targeting strategy was developed by the authors, and the mouse generation service was performed by Shanghai Model Organisms, Shanghai, China, by co-injecting into zygotes the Cas9 protein, single guide RNA (sgRNA), and a single-stranded mutant oligo DNA [70]. The donor DNA template contains a c.1151C>A (p.Ser384*) mutation in *Akap11* (ENSMUST00000123853), which was homologous to the *AKAP11* PTV chr13-42299894C>A in the BipEx database [11] that encodes a c.1148C>A (p.Ser383*) mutation (ENST00000025301). Founder (F0) mice were screened by polymerase chain reaction (PCR) amplification of the targeted locus followed by Sanger sequencing. One correctly targeted founder was bred with C57BL/6J wild-type mice to establish the line. Experimental animals were genotyped by PCR followed by Sanger sequencing or by real-time PCR. All animals used were derived from one heterozygous F2 male mouse. Experiments were conducted using mice from F4 and later generations. Genotyping primers for the mutation are provided in Supplementary Table 1C. *sgRNA off-target prediction*

To predict potential off-target effects, the guide RNA (gRNA) sequence was analyzed using Synthego CRISPR design tool (https://www.synthego.com/products/bioinformatics/crispr-design-tool) and the top ranked five hits were selected for screening. Off-target (OT) primers (OT-F/OT-R) were designed to amplify 400-600 bp regions spanning each potential off-target site using Tag master mix (Vazyme, Nanjing, China). Single PCR product was confirmed for each primer pair. PCR products were Sanger sequenced to confirm the absence of indels at the predicted off-target sites. PCR primers for off-target sites are provided in Supplementary Table 1C.

#### Animal housing and handling

The animals were handled in accordance with protocols approved by the Johns Hopkins Animal Care and Use Committee and with the National Institutes of Health guidelines. For all experiments except behavioral assays and primary neuronal culture experiments, male mice were housed in groups of three to five per cage in a standard 12-h light/12-h dark cycle and were provided unlimited water and food. Tissues were harvested from 12-week-old mice during the dark phase. Only male mice were used unless otherwise indicated. Primary neuronal cultures were prepared from embryonic day 16.5 (E16.5) mice of both sexes.

### Behavioral analysis

#### Animal housing and stress induction

For behavioral experiments, 10 *Akap11^+/S384*^* and 10 *Akap11^+/+^* male mice were housed in groups of four with balanced genotypes per cage. Mice were transferred to reverse light/dark cycle at least two weeks prior to behavioral testing. Baseline behavioral tests were performed at 12 weeks of age. Mice were then housed singly without environmental enrichment for three weeks, followed by post-stress behavioral testing. All behavioral testing was conducted during the dark phase. The testing chamber and room were kept dark with red LED lighting in the room. Before each test, mice were habituated to the behavioral testing room for 30 min to 1 h.

#### Open field test

Each test mouse was placed in the center of an open field testing chamber equipped with infrared sensors (San Diego Instruments, San Diego, CA, USA). Horizontal and vertical movements were recorded for 2 h in the dark. Infrared beam breaks were recorded in 5-min bins.

#### Elevated plus maze

Each test mouse was placed in the center zone of the elevated plus maze, and allowed to explore the maze for 5 min. Movements in the open arms, closed arms, and center zone of the maze were tracked and recorded using the ANY-maze software (version 7.65).

#### Social interaction test

The three-chamber social interaction test was carried out as previously described with modifications [71]. During the habituation phase, the mouse chamber and toy chamber both contained empty cups. The test mouse was placed into the center zone of the apparatus and allowed to freely explore all three chambers for 6 min. During the test phase, an unfamiliar, age-matched *Akap11^+/+^* male mouse was placed under the cup in the mouse chamber, and a toy mouse was placed under the cup in the toy chamber. The test mouse started from the center zone and was allowed to explore the social and non-social stimuli for 10 min. Behavior was video-recorded using ANY-maze. An experimenter blinded to the genotype of the test mouse manually scored the following measurements: (a) time spent in the mouse chamber (*T_mouse chamber_*) or toy chamber (*T_toy chamber_*), defined as the test mouse having at least 50% of its body length being inside the respective chamber; and (b) time spent interacting with the social stimulus (*T_mouse_*) or non-social stimulus (*T_toy_*), as defined by Rein et al. [71]. The chamber-based sociability index was computed as *SI_chamber_* = *T_mouse chamber_* / *T_toy chamber_*. The interaction-based sociability index was computed as *SI_interaction_* = *T_mouse_* / *T_toy_*.

### Tissue harvesting for reverse transcription-quantitative PCR, full-length RNA sequencing, and single-nucleus RNA sequencing

Mice were anesthetized with isoflurane and transcardially perfused with 10 mL ice-cold nuclease-free 1X phosphate buffered saline (PBS; Invitrogen, Waltham, MA, USA). For reverse transcription-quantitative PCR (RT-qPCR) and long-read RNA sequencing, the portion of the brain rostral to the optic chiasm was sectioned into 1.5 mm coronal slices using a prechilled brain matrix. The somatosensory cortex was rapidly dissected from those sections and snap-frozen on dry ice. For single-nucleus RNA sequencing, the neocortex was rapidly isolated from subcortical structures. The sensory cortical areas (somatosensory, visual, and auditory cortices), the medial area (retrosplenial area), and lateral areas (the temporal association, ectorhinal, and perirhinal areas) were collected and snap-frozen on dry ice.

### RT-qPCR

Somatosensory cortices from three *Akap11^+/S384*^* and four *Akap11^+/+^* mice (male, 12 weeks old) were used for RT-qPCR. Tissues were homogenized in TRIzol Reagent (Thermo Fisher Scientific, Waltham, MA, USA) using a BeadBug tissue homogenizer (Benchmark Scientific, Edison, NJ, USA). RNA was then purified using the Direct-zol RNA Miniprep kit (Zymo Research, Irvine, CA, USA) following the manufacturer’s instructions. Total RNA was reverse transcribed using HiScript IV RT SuperMix for qPCR (+gDNA wiper) kit (Vazyme). qPCR was performed using Taq Pro Universal SYBR qPCR Master Mix (Vazyme) in the QuantStudio 12K Flex system (Thermo Fisher Scientific). RT-qPCR primers for *Akap11* and the reference genes are provided in Supplementary Table 1C.

### Full-length RNA sequencing library preparation and sequencing run

Full-length RNA sequencing was performed using somatosensory cortices from three homozygous mutants (*Akap11^S384*/S384*^*) and three control mice (*Akap11^+/+^*) (male, 12 weeks old). RNA extraction, library preparation, and sequencing were performed by Admera Health (South Plainfield, NJ, USA). Briefly, total RNA was extracted using the RNeasy Kits (Qiagen, Hilden, Germany) following the manufacturer’s instructions. RNA sample quality was assessed with the High Sensitivity RNA ScreenTape Assay on the Tapestation (Agilent Technologies, Santa Clara, CA, USA) and quantified by the AccuBlue Broad Range RNA Quantitation assay (Biotium, Fremont, CA, USA).

Full-length cDNA libraries were generated using the Kinnex full-length RNA kit (PacBio, Menlo Park, CA, USA) following the manufacturer’s protocol. Briefly, 450 ng of RNA was used for cDNA synthesis using the Iso-Seq Express 2.0 Kit (PacBio). For each genotype, 18.3 ng of barcoded cDNA from each sample was pooled for eight parallel Kinnex PCR, followed by Kinnex array formation, nuclease treatment, and SMRTbell library preparation. Quality of the final library was assessed using the Genomic DNA ScreenTape Assay (Agilent Technologies), and quantity was measured using Qubit 2.0 dsDNA HS assay Kit (ThermoFisher). Sequencing was performed on one SMRTCell on the Revio system (PacBio).

### Single-nucleus RNA sequencing

Two *Akap11^+/S384*^* mice and three *Akap11^+/+^* littermates (male, 12 weeks old) were used for single-nucleus RNA sequencing (snRNA-seq). Tissue dissociation, nuclei isolation, library preparation, and sequencing were performed by Admera Health. Briefly, tissue dissociation and nuclei isolation was performed with nuclei extraction buffer (Miltenyi Biotec, Bergisch Gladbach, Germany) and the gentleMACS Dissociator (Miltenyi Biotec) following the manufacturers’ recommendation. Nuclei were washed and resuspended in PBS with 0.04% BSA. Nuclei concentration and quality were assessed on the Cellometer Auto 2000 (Nexcelom, Lawrence, MA, USA) with Acridine Orange and Propidium iodide stain. Single-nucleus 3’ Gene Expression libraries were generated using the Chromium GEM-X Single Cell 3’ Kit v4 (10x Genomics, Pleasanton, CA, USA) following the manufacturer’s instructions. Briefly, 30000 nuclei per sample were loaded on the Chromium X (10x Genomics) for cell partitioning and Gel Beads-in-emulsion (GEM) generation. Following GEM generation, reverse transcription was performed, and cDNA was pooled, purified, and amplified. Libraries were constructed using 13 cycles in Sample Index PCR. Quality of cDNAs and libraries was assessed using High Sensitivity D5000 ScreenTape (Agilent Technologies) and quantified with the Qubit 2.0 dsDNA HS assay Kit (ThermoFisher). Equimolar pooling of libraries was performed based on quality control (QC) values. Libraries were sequenced on NovaSeq X Plus (Illumina, San Diego, CA, USA) with PE-150 read length configuration.

### Immunoblotting

Six *Akap11^+/S384*^* and six *Akap11^+/+^* mice (male, 12 weeks old) were used for immunoblotting. Mice were anesthetized with 300-500 μL isoflurane. The brain was removed from the skull, and the region rostral to the optic chiasm was sliced into 1.5 mm coronal sections using a prechilled mouse brain matrix. The somatosensory cortex, prefrontal cortex, and striatum were rapidly dissected from these sections under a dissection microscope, and the hippocampus was dissected from the remaining caudal tissue. Tissues were snap-frozen on dry ice.

Tissues were homogenized using a plastic pestle (Genesee Scientific, San Diego, CA, USA) in RIPA Buffer (Sigma-Alrich, St. Louis, MO, USA) containing protease inhibitors (Roche Diagnostics, Mannheim, Germany), phosphatase inhibitor cocktail 2 (Sigma-Aldrich) and phosphatase inhibitor cocktail 3 (Sigma-Aldrich). Protein concentration was measured using the Pierce BCA Protein Assay Kit (Thermo Fisher Scientific).

For each sample, 30 µg of total protein was loaded per lane on 4-20% Bis-Tris gels (GenScript, Piscataway, NJ, USA) and separated by eletrophoresis in Tris-MOPS-SDS running buffer (GenScript). Proteins were transferred to a 0.45 µm low-fluorescence PVDF membrane (Azure Biosystems, Dublin, CA, USA). For fluorescent immunoblots, membrane blocking and antibody incubation were done using the Azure Fluorescent Blot Blocking Buffer (Azure Biosystems). For combined chemiluminescent and fluorescent immunoblots, the Rockland Blocking Buffer (Rockland Immunochemicals, Limerick, PA, USA) was used. Primary antibodies were incubated overnight at 4 °C with gentle agitation. Secondary antibodies were incubated for 1-2 h at room temperature with gentle agitation. Membranes were imaged and quantified using the LI-COR Odyssey Fc Imager and Image Studio Software (version 5.2.5; LI-COR Biosciences, Lincoln, NE, USA).

Immunoblotting was performed using the following primary antibodies: rabbit anti-AKAP11 (CST 77313, 1:1000), rabbit anti-PKA RIα (CST 5675, 1:1000), sheep anti-PKA RIβ (R&D Systems

AF4177, 1:1000), rabbit anti-PKA Cα (CST 4782, 1:1000), rabbit anti-PKA Cβ (Proteintech 12232-1-AP, 1:600), mouse anti-PKA RIIα (BD Biosciences 612242, 1:1000), mouse anti-PKA RIIβ (BD Biosciences 610625, 1:1000), mouse anti-GAPDH (Santa Cruz Biotechnology sc-32233, 1:5000-1:3000), and rabbit anti-GAPDH (CST 2118, 1:5000-1:3000). The following secondary antibodies were used: goat anti-Rabbit IgG (HRP) (CST 7074, 1:5000), donkey anti-Sheep IgG (HRP) (R&D Systems HAF016, 1:1000), goat anti-Mouse IgG (IR700) (Azure Biosystems AC2129, 1:5000), and goat anti-Rabbit IgG (IR800) (Azure Biosystems AC2134, 1:5000).

### Immunocytochemistry

Cultured neurons transfected with pCAG-GFP between DIV4 and DIV6 were fixed between DIV21 and DIV22 for immunocytochemistry. Neurons were fixed with 4% paraformaldehyde (PFA) (Electron Microscopy Sciences, Hatfield, PA, USA) in PBS for 20 min at room temperature and permeabilized with 0.1% Triton X-100 (Sigma-Alrich) in PBS for 15 min. Cells were blocked in 5% bovine serum albumin (BSA) (Sigma-Alrich) in PBS for 1 h at room temperature, and then incubated overnight at 4 °C with primary antibodies diluted in 5% BSA. Secondary antibodies were diluted in 5% BSA and incubated for 1 h at room temperature.

Coverslips were counterstained with 0.5 µg/mL 4’,6-diamidino-2-phenylindole (DAPI) (Sigma-Alrich) for 10 min at room temperature before mounting to microscope slides (Thermo Fisher Scientific) with Aqua-Poly/Mount medium (Polysciences, Inc., Warrington, PA, USA). For immunocytochemistry, the following primary antibodies were used: rabbit anti-PKA RIα (CST 5675, 1:1000), rabbit anti-MAP2 (CST 8707, 1:1000), and mouse anti-Tau-1 (Sigma-Aldrich MAB3420, 1:1000). The following secondary antibodies were used: donkey anti-Rabbit IgG (H+L) (Alexa Fluor 488) (Thermo Fisher Scientific A-21206, 1:1000), and donkey anti-Mouse IgG (H+L) (Alexa Fluor 555) (Thermo Fisher Scientific A-31570, 1:1000).

### Proteomics and phosphoproteomics

#### Processing of tissues

Five *Akap11^+/S384*^* and four *Akap11^+/+^* mice were used for quantitative bulk proteomics and phosphoproteomics. Mice were euthanized by cervical dislocation. The olfactory bulb was removed, and the neocortex was rapidly isolated from subcortical structures using a razor blade. After heat stabilization on the Stabilizor T1 (Denator AB, Uppsala, Sweden) to inactivate enzymes, tissues were snap-frozen on dry ice.

#### Sample preparation, Tandem mass tag labeling, and fractionation

Proteomics sample preparation and tandem mass tag (TMT)-based liquid chromatography-tandem mass spectrometry (LC-MS/MS) were performed by the Johns Hopkins Center for Proteomics Discovery. Sample preparation was conducted as described previously with minor modifications [72, 73]. Tissues were lysed in 8 M urea/50 mM triethylammonium bicarbonate with protease/phosphatase inhibitors and sonicated on ice. Proteins (100 μg/sample) were reduced with 10 mM tris(2-carboxyethyl)phosphine, alkylated with 40 mM 2-chloroacetamide, digested with Lys-C (1:50), diluted to approximately 2 M urea, and digested overnight with trypsin (1:50, 37 °C) [74]. Peptides were acidified, C18-desalted, labeled with TMTpro 18-plex reagents (Thermo Fisher Scientific) for 1 h at RT before quenching with Tris and pooling [73]. Samples were fractionated by basic pH reversed-phase liquid chromatography (bRPLC) on an Agilent 1260 HPLC/Extend-C18 column (Agilent Technologies) using a 90 min gradient at 0.3 mL/min [72]. 10% of the fractions was used for global proteomics and 90% for Fe(III)-IMAC phosphopeptide enrichment [75].

#### LC-MS/MS and database searching

Peptides were analyzed using Vanquish NEO UHPLC with Orbitrap Exploris 480 (Thermo Fisher Scientific) with a 120 min gradient at 300 nL/min. MS1 spectra were acquired at 120,000 resolution (m/z 375-1500), followed by data-dependent acquisition (3 s cycle, 0.7 Da isolation, 60 s exclusion) and higher-energy collisional dissociation (normalized collision energy of 36); MS/MS spectra were acquired at 30,000 resolution using TurboTMTPro. Spectra were searched with SEQUEST HT in Proteome Discoverer (version 3.2.0.450, Thermo Fisher Scientific) against the UniProt Mus musculus database [76]. Searches allowed two missed tryptic cleavages, ≥ 6-aa peptides, and 10 ppm/0.02 Da mass tolerances. Carbamidomethylation and TMTpro labeling were fixed; oxidation, deamidation, and, for phosphoproteomics, Ser/Thr/Tyr phosphorylation were variable. False discovery rate (FDR) control was applied at 1% at both peptide-spectrum match (PSM) and protein levels using the Percolator node implemented in Proteome Discoverer. Phosphosite localization confidence was assessed using the ptmRS node. Reporter ion quantification was performed using MS2-based TMT quantification with a reporter ion integration tolerance of 20 ppm. Protein grouping was performed according to the principle of strict parsimony.

### DNA constructs

The PKA sensor pcDNA3.1(+)-ExRai-AKAR2 was a gift from Jin Zhang (Addgene plasmid #161753; RRID: Addgene_161753). To generate the pAAV-Camk2a-ExRai-AKAR2-mScarlet plasmid, the pCamk2a-PKA sensor-P2A-mScarlet sequence was gene-synthesized by Genscript and cloned into the Xba1/EcoR1-linearized pAAV-hSyn-EGFP backbone. pAAV-hSyn-EGFP was a gift from Bryan Roth (Addgene plasmid #50465; RRID: Addgene_50465). For sparse labeling in neuronal cultures, pCAG-GFP was a gift from Connie Cepko (Addgene plasmid #11150; RRID: Addgene_11150).

### Primary neuronal cultures and transfection

Primary dissociated cultures were prepared from cortices of E16.5 embryos. Cortices were dissected in Dulbecco’s Modified Eagle’s Medium (DMEM) (Corning, Corning, NY, USA) supplemented with 1X GlutaMAX (Thermo Fisher Scientific), 0.5% penicillin/streptomycin (Corning), and 10% FBS (Corning). After washing with PBS, tissues from individual embryos were cut into small pieces, and dissociated with TrypLE (Thermo Fisher Scientific) supplemented with DNase I (Roche) at 37 °C. Tissues were then washed with culture medium consisting of Neurobasal medium (Thermo Fisher Scientific) supplemented with B-27 (Thermo Fisher Scientific), GlutaMAX, and 0.5% penicillin/streptomycin, followed by trituration.

Dissociated cells were plated on poly-L-ornithine (Sigma-Alrich) coated coverslips or imaging dishes. Cells were seeded at 1.5 x 10^5^ cells per 12 mm coverslip (Electron Microscopy Sciences) or 4.5 x 10^5^ cells per 35 mm imaging dish (Cellvis, Mountain View, CA, USA). For morphological analysis of nascent neurites, cells were seeded at 1.0 x 10^3^ cells per 12 mm coverslip. Cultures were maintained at 37 °C in a humidified incubator with 5% CO_2_. *Post hoc* genotyping for *Akap11* mutation and sex was performed using DNA extracted from the tail of each embryo. Genotyping primers are provided in Supplementary Table 1C.

Transient transfection of the PKA sensor was performed between days *in vitro* 5 and 6 (DIV5-DIV6) using Lipofectamine 2000 (Invitrogen). Cells in each imaging dish were transfected with 1 µg of plasmid DNA mixed with 2 µL Lipofectamine, and incubated in Neurobasal medium for 4 hours, before returning to the conditioned culture medium. Transient transfection with pCAG-GFP was performed at DIV4-6 at a 1:1 DNA-to-Lipofectamine 2000 ratio, with 200 ng DNA per 12-mm coverslip.

### Time-lapse imaging of PKA activity

Cultured cortical neurons transfected a PKA sensor were imaged between DIV21 and DIV22 on the Nikon Ti2 widefield microscope (Nikon Instruments, Tokyo, Japan) equipped with a CFI Apochromat TIRF 60X/1.49 NA oil objective (Nikon Instruments), an ORCA-Fusion Digital CMOS camera C14440-20UP (Hamamatsu Photonics, Hamamatsu, Japan), and controlled with the NIS-Elements AR software (version 6.10.02, Nikon). Dual GFP excitation imaging was performed using the 405 nm excitation (10% laser power, 500 ms exposure time) and 488 nm excitation (2.5% laser power, 500 ms exposure time) and a 525/50 nm emission filter. Red fluorescence from mScarlet was imaged using a 561 nm excitation (5% power, 50 ms exposure time) and a 595/44 nm emission filter.

For live-cell imaging, baseline images were acquired every 1 min for 5 min. Forskolin (25 µM; Cell Signaling Technology, Danvers, MA, USA) plus rolipram (2 µM; Sigma-Aldrich) or H-89 (25 µM; Cell Signaling Technology) was then added to the imaging dish. Cells were incubated in the drugs for 3 min before resumption of imaging at 1 min intervals for 15 min.

### Statistical analysis

Behavioral data, physical measurements, immunoblotting, RT-qPCR, immunocytochemistry, and live-cell PKA activity imaging data were analyzed using R (version 4.3.0) running in a Debian GNU/Linux 11.7 container image. For behavioral analyses, the interquartile range (IQR) was calculated for each genotype, and data points falling 1.5 x IQR below the first quantile (Q1) or above the third quantile (Q3) were removed as outliers. For physical measurements, no data points were excluded. A linear regression with genotype and litter as fixed effects was fit to rank-transformed data, and *post-hoc* pairwise comparison was performed using Tukey’s test. For immunoblotting, RT-qPCR, immunocytochemistry, morphometric analysis, and live-cell imaging of PKA activity, no data points were excluded. Mann-Whitney U test was used to determine the difference between the two genotypes.

### Processing and analysis of full-length RNA sequencing data

Processing and analysis of the full-length RNA sequencing data followed the *IsoSeq* workflow (version 4.3.0) with Python 2.7.15. HiFi reads were segmented using *Skera* (version 1.4.0) with MAS-Seq Adapter v3. Demultiplexing and sample barcode identification were performed using *lima* (version 2.13.0) with the Iso-Seq v2 12-plex barcoding primers in the Iso-Seq mode with peak guessing (--isoseq --peek-guess). Poly(A) tails were trimmed and concatemers removed using *isoseq refine* (--require-polya) with the default minimum poly(A) tail length of 20 bp. The full-length non-concatemer (FLNC) reads were then clustered using *isoseq cluster2*, keeping only isoforms with at least two FLNC reads. Clustered isoforms were then aligned to the GRCm39/mm39 mouse reference genome using the *minimap2* wrapper *pbmm2* (version 1.17.0; --preset ISOSEQ --sort). Redundant isoforms were collapsed using *isoseq collapse* (--do-not-collapse-extra-5exons --max-fuzzy-junction 10 --max-5p-diff 100 --max-3p-diff 100).

Nonredundant isoforms were classified using *pbpigeon* (version 1.4.0) with the *pigeon classify* command, the mouse reference genome (GRCm39/mm39), the sorted and indexed GENCODE vM33 genome annotation, and a curated list of poly(A) sites (PacBio). Outputs were filtered using *pigeon filter* (--min-cov=3 --mono-exon) to remove low-confidence and mono-exonic isoforms. For downstream QC, we further removed isoforms with (a) a junction predicted to be template-switching artifact (“RTS_stage”=TRUE), (b) > 80% adenine nucleotides within the 20 bp window immediately downstream of a transcript’s termination site (TTS) (“perc_A_downstream_TTS” > 80), or (c) total number of counts < 3. The remaining isoforms were normalized to transcripts per million (TPM). Isoforms for *Akap11* were visualized using UCSC Genome Browser on GRCm39/mm39 (https://genome.ucsc.edu/).

### Processing and analysis of proteomic and phosphoproteomic data

#### Processing and analysis of proteomics data

Proteomic data were analyzed in R (version 4.5.3) running in an Ubuntu 20.04.6 LTS container image. Proteomic data was processed as *Msnbase* objects (version 2.34.1), and proteins that were classified as contaminants were removed. Protein abundance for each sample was log_2_ transformed and normalized using the cyclic loess method via *normalizeBetweenArrays* (method=“cyclicloess”) in the package *limma* (version 3.64.3). Differential abundance analysis was performed by fitting a linear model to the data via *limma::lmFit*, followed by empirical Bayes moderation of the standard errors via *limma::eBayes* (trend=TRUE, robust=TRUE).

#### Processing and analysis of phosphoproteomic data

Phosphosite data was processed in *Msnbase* format, log_2_ transformed, and cyclic loess-normalized to generate the phosphosite abundance data. To estimate the phosphorylation rate, bulk proteome and phosphoproteome abundance values were first normalized within each sample to the channel median. Phosphosite abundance was then normalized to the corresponding bulk proteome abundance, and log_2_-transformed. The resulting phosphorylation rate values were further cyclic loess-normalized. Differential phosphosite abundance and phosphorylation rate analyses were performed using *limma::lmFit* and *limma::eBayes*.

#### Kinase activity analysis

Kinase-substrate annotations were obtained from the PTMsigDB collection (version 2.0.0) [77], retrieved on April 05, 2026, and from curated RoKAI annotations (version 2.3.0) [49], accessed on April 05, 2026. The RoKAI annotations were based on PhosphositePlus [47] and Signor [48]. The “DE scores” for phosphosites were used as inputs for kinase activity inference, which was calculated as −log_10_*P* x sign(log_2_ fold change). Following the RoKAI method [49], activity score for a kinase was computed as the mean DE scores of its substrates and the associated *P* value was estimated using a *Z* test. We also applied an alternative kinase activity test based on Alvarez et al. [78] via the *viper* package (version 1.42.0) using PTMsigDB annotations, which employs analytic rank-based enrichment analysis to estimate the enrichment scores, and permutation test to estimate *P* values.

### Processing and analysis of snRNA-seq data

#### snRNA-seq data processing

Sequenced reads were aligned to a preprocessed GRCm39/mm39 reference genome (GRCm39-2024-A; 10x Genomics) using Cell Ranger (version 9.0.1) with the *cellranger count* pipeline (--create-bam true, --include-introns=true, and default values for other parameters). The count matrix was processed as *SingleCellExperiment* objects (version 1.24.0). Downstream QC and analyses were performed in R (version 4.3.0) running in a Debian GNU/Linux 11.7 container image.

#### Quality control

Potential doublets were identified using the *scDblFinder* package (version 1.14.0) to compute the density-based doublet scores with *computeDoubletDensity* (d=50), followed by thresholding with *doubletThresholding* (dbr=0.10, method=“griffiths”, returnType=“call”, p=0.01). Next, we filtered cells on a per-sample basis with following criteria: (a) total unique molecular identifiers (UMIs) < 1000 or > 5 median absolute deviations (MADs); (b) uniquely detected genes < 750 or > 5 MADs; or (c) percentage of UMIs mapped to mitochondrial genes > 5 MADs. Finally, we removed genes that were expressed fewer than 10 copies in all samples.

#### Clustering and cell-type annotation

We applied the following pipeline to cluster cells and assign cell type identity at multiple resolutions:

1. Normalization. Gene expression was normalized using *normalizeCounts* in the *scuttle* package (version 1.12.0). Size factors were calculated from each cell’s total UMI counts, scaled by the geometric mean of all cells.
2. Feature selection. Mitochondrial and ribosomal genes were excluded. Highly variable genes were identified using variance stabilizing transformation by fitting a LOESS regression to the variance and mean in log scale using *loess* from the *stats* package (version 4.3.0). The expected variance was clipped at 0.3. Standardized variances were transformed into *Z* scores, and right-tailed *P* values were calculated and adjusted by FDR correction. Highly variable genes were selected by FDR < 0.50, or as the top 3000 genes with the highest standardized variance if fewer than 3000 genes were available by the first criterion.
3. Dimensionality reduction. Principal component analysis (PCA) was performed using the highly variable genes. PCA was computed with *calculatePCA* from the *scater* package (version 1.28.0). For each dataset, to determine the optimal number of components, an initial PCA was performed using 50 components, and the cumulative variance explained was computed. The optimal number of components was defined as the largest number of PCs for which the cumulative variance explained did not exceed 80% in the initial PCA, except for the clustering of cells into broad cell types (Glutamatergic, GABAergic, and non-neuronal), where 40 PCs were used. Uniform manifold approximation and projection (UMAP) was computed based on the PCs using *calculateUMAP* in *scater* with the optimized number of PCA dimension (“n_dimred”). The number of nearest neighbors (“n_neighbors”) was set at 20.
4. Shared nearest-neighbor (SNN) graph construction. SNN graphs were constructed using *buildSNNGraph* in the *scran* package (version 1.28.2) with 20 nearest neighbors, and the rank-based weighting scheme.
5. Clustering. The SNN graph was partitioned using the Leiden algorithm at optimal resolutions, implemented by *cluster_leiden* in the *igraph* package (version 1.5.0.1; objective_function=“CPM”, n_iterations=5). To assess robustness of the clusters, we repeated steps 2-5 with stratified and balanced sampling of the input data such that each cell was re-clustered 50 times (for the initial clustering) or 100 times (for subclustering). Clusters generated across resampling iterations were matched to clusters from a reference iteration using pairwise Adjusted Rand Index. The probability of a cell being assigned into the same reference cluster was computed.
6. Cell type assignment. Clusters were annotated in accordance with the mouse cortical cell-type taxonomy proposed by Yao et al. [30] with modifications. A set of reference marker genes were curated by integrating: (a) canonical cell-type markers reported in the literature [29, 30]; (b) a reanalysis of the Allen Institute Whole Cortex and Hippocampus 10x Genomics dataset, retrieved from https://brain-map.org/our-research/cell-types-taxonomies/cell-types-database-rna-seq-data/mouse-whole-cortex-and-hippocampus-10x. To obtain reference marker genes in the Whole Cortex and Hippocampus dataset, we randomly subsampled up to 1211 cells per subclass in this dataset. Cells assigned to the hippocampal formation were excluded, as were cells belonging to subclasses primarily identified in the hippocampal formation (“ENT”, “HIP”, and “PAR-POST-PRE-SUB-ProS” as “region_label”). Cells from the “Meis2” subclass were also excluded as this subclass contained only 20 cells. Pairwise Wilcoxon tests were performed using *findMarkers* in the *scran* package (test.type=“wilcox”, pval.type=“all”, block=“dataset”) to identify markers for the following cell types:

i. Broad cell types: Glutamatergic neuron, GABAergic neuron, and the non-neuronal cell types (“Astro”, “Endo”, “Micro-PVM”, “Oligo”, “SMC-Peri”, and “VLMC”).
ii. Layer-specific glutamatergic cell types: “L2/3 IT CTX”, “L2/3 IT ENT”, “L2/3 IT PPP”, “L2 IT RSP-ACA”, “L4 RSP-ACA”, “L4/5 IT CTX”, “L5 IT CTX”, “L5/6 IT TPE-ENT”, “L5 PT CTX”, “L5/6 NP CTX”, “L6 IT CTX”, “L6 CT CTX”, “L6b CTX”, and “Car3”.
iii. Caudal ganglionic eminence (CGE)-derived GABAergic cell types: “Lamp5”, “Sncg”, and “Vip”.
iv. Medial ganglionic eminence (MGE)-derived GABAergic cell types: “Pvalb”, “Sst”, and “Sst Chodl”.

Initial cell type assignment was performed using the cell-type specificity score described by Ianevski et al. [79], with the curated set of reference markers. Assignment was then corroborated with cluster-enriched genes (obtained by pairwise Wilcoxon tests between cells within and outside a cluster). For glutamatergic cell types, annotations were additionally corroborated with the laminar expression patterns of cluster-enriched genes in the Allen Mouse Brain in situ hybridization database (https://mouse.brain-map.org/).

#### Clustering and subclustering workflow

Initial clustering was performed on the full data to identify the broad cell-type classes: Pan-neuron, Oligodendrocyte, OPC, Endothelial cell, VSMC, Astrocyte, Microglia, Pericyte, and Meningeal cell. Cells in the pan-neuronal clusters were further clustered into the “Glutamatergic”, “GABAergic”, and “mixed/other” subclasses. Glutamatergic neurons were subclustered into 13 layer- and/or region-specific cell types, while GABAergic neurons were subclustered into 10 cell types. Neurons in the “mixed/other” subclass were removed from downstream analysis. Initial clustering and subclustering were performed using a range of resolutions in the Leiden clustering, and the optimal resolution was chosen as one that enabled identification of the largest number of known cell types described in prior mouse cortical taxonomies [29, 30].

Each glutamatergic, GABAergic, and non-neuronal cell types were then further clustered into subtypes by applying the clustering pipeline at a range of finer resolutions. To select the optimal resolution for glutamatergic cell types, we performed pairwise Wilcoxon tests between resulting clusters for each gene via *scran::pairwiseWilcox*, and computed a *deScore* as the mean - log_10_FDR. We also evaluated cluster robustness at each resolution by computing the proportion of cells assigned to the same reference cluster in at least 75% of resampling iterations, referred to as the “valid cell ratio”. Optimal resolutions were chosen to yield comparable minimum *deScore* values across glutamatergic cell types (average: 4.83, min: 3.78, max: 7.29), while maintaining high valid cell ratio (average: 98.65%, min: 90.15%, max: 100%). For GABAergic cell types, optimal subclustering resolutions were chosen to enable recovery of previously described interneurons subtypes [29]. For non-neuronal cell types, the Oligodendrocyte, OPC, Endothelial cell, Astrocyte and Pericyte were further clustered. The optimal resolution was chosen at the point where both the minimum *deScore* and the number of clusters reached a plateau.

#### Differential expression analysis

Evaluation of cell type and subtype composition between genotypes was performed using *propeller* in the *speckle* package (version 1.0.0; robust=TRUE, transform=“logit”).

For differential expression (DE) analysis, samples containing fewer than 10 cells from a given cluster were excluded for that cluster. Clusters with fewer than two samples per genotype after this filtering step were excluded from DE analysis. For pseudobulk DE analysis, gene-level total counts were aggregated within each sample and cluster using *scater::aggregateAcrossCells* (statistics=“sum”). Lowly expressed genes were filtered using *filterByExpr* in the *edgeR* package (version 4.0.16) with default parameters. Library normalization factors were computed using the relative log expression (RLE) method with *edgeR::calcNormFactors* (method=“RLE”). Gene expression was modeled by gene-wise quasi-likelihood negative binomial generalized log-linear models implemented in *edgeR::glmQLFit* (robust=TRUE) and *edgeR::glmQLFTest*. Empirical Bayes moderation was applied to the quasi-likelihood dispersions, and *P* values were estimated via F-tests. Adjusted *P* values were computed using the FDR method. Alternative pseudobulk DE analyses were performed using *DESeq2* (version 1.40.2) and *limma* (version 3.58.1).

To perform generalized linear mixed model (GLMM)-based DE test on UMI counts from individual cells, we filtered samples and clusters using the same criteria as for the pseudobulk analysis, and computed the normalization factor with the RLE method, scaled by the library size. For each gene, a Poisson-gamma mixed model was fit using the *nebula* function in the *nebula* package (version 1.5.5; model=“PMM”, cpc=0.005, mincp=20). Adjusted *P* values were computed using the FDR method.

### Pathway enrichment analysis

#### Pathway databases

Mouse Gene Ontology (GO) annotation databases were retrieved from the *org.Mm.eg.db* package (version 3.18.0) using *clusterProfiler* (version 4.10.1) on September 10, 2025. Three classes of GO sub-ontology were used: biological process (GO:BP), cellular component (GO:CC), and molecular function (GO:MF). Mouse Reactome pathway database was retrieved via the *ReactomePA* package (version 1.46.0) on September 10, 2025. The expert-curated synaptic gene-set database SynGO (version 20231201) was retrieved from the SynGO portal (https://www.syngoportal.org). Gene sets associated with psychiatric disorders were collected from published studies [5, 10, 11, 13, 24, 80–92]. For GWAS-based risk genes, we selected gene sets prioritized by gene-based association analyses or by integration of multiple lines of evidence. Supplementary Table 3G summarizes gene-set processing procedures applied to each dataset, including the data source, prioritization method, and *P* value cutoffs if applicable. We also retrieved 75 annotated gene sets from DisGeNET (2026 update) under the Medical Subject Headings (MeSH) category F03, “Mental Disorders” (accessed July 06, 2026).

#### Overrepresentation analysis

Overrepresentation analysis (ORA) was performed using *fora* in the *fgsea* package (version 1.26.0; minSize=5, maxSize=21955). *P* values were estimated via hypergeometric tests and adjusted by the FDR method. For snRNA-seq data, the background gene set was defined as all genes meeting the minimum expression criteria in our data. Upregulated and downregulated genes were analyzed separately.

#### Gene set enrichment analysis

Genes were ranked by -log_10_*P* x sign(log_2_ fold change). Gene set enrichment analysis (GSEA) was performed using *fgsea::fgseaMultilevel* (scoreType=“std”, maxSize=500, eps=0, nPermSimple=10000). The minimum gene-set size was set to 10 for snRNA-seq data, and 15 for proteomic data. *P* values were estimated via a permutation test with 10000 permutations and adjusted by the FDR method. To robustly estimate enrichment, the GSEA test was repeated 20 times. Pathways were considered robustly enriched if they reached significance (FDR ≤ 0.05) in at least 70% of runs. For snRNA-seq data, to ensure statistical reliability of the permutation tests, we restricted GSEA to cell types or subtypes in which at least 5000 genes met the minimum expression requirement for DE analysis. For proteomic and phosphoproteomic data, UniProt accession of proteins were mapped to MGI gene symbols using *biomaRt* (version 2.58.2). When multiple UniProt accessions mapped to the same MGI symbol, the Uniprot accession with the largest absolute ranking score was retained. For phosphosite abundance and phosphorylation rate data, GSEA was performed on the genes corresponding to the master proteins. When multiple phosphosites were mapped to the same gene, the entry with the largest absolute ranking score was used.

#### Semantic summary of significant pathways

To facilitate interpretation of ORA and GSEA results, significant pathways were clustered based on semantic similarity using the *rrvgo* package (version 1.14.2). Similarity between pathways were computed using *rrvgo::calculateSimMatrix* (orgdb=“org.Mm.eg.db”, method=“Lin”), and pathways were grouped by *rrvgo::reduceSimMatrix* (orgdb=“org.Mm.eg.db”, scores=“uniqueness”). Clustering of the pathways were further manually curated by comparison with GO ancestor charts retrieved from QuickGO (https://www.ebi.ac.uk/QuickGO/; retrieved September 10, 2025) to (a) merge related clusters spanning different GO sub-ontologies, and (b) split clusters when the automated clustering was inconsistent with GO ancestor relationships.

### Quantification of nascent neurite morphology

Cultured neurons at DIV6 were imaged on the Axio Observer Z1 widefield microscope (Zeiss, Oberkochen, Germany) equipped with an Axiocam705 and a 40x/0.95 NA objective (Plan-Apochromat 40x/0.95 Corr M27; Zeiss). Tiled images were acquired at a resolution of 11.59 pixels per µm at a single depth. Images were acquired using identical exposure time and binning mode across cells. From each coverslip, cells showing a single Tau-1-positive neurite and multiple MAP2-positive neurites were randomly selected for imaging. Neurons were manually traced in Neurolucida (version 2025.1.2; MBF Bioscience LLC, Williston, VT, USA) by an experimenter blinded to the neuronal genotype. Morphometric measurements were calculated using Neurolucida Explorer (version 2024.1.1; MBF Bioscience LLC) and analyzed in R (version 4.3.0) using the Mann-Whitney U test.

### Quantification of immunocytochemistry

Neurons were imaged on the Axio Observer Z1 microscope with the LSM 800 confocal laser scanning module (Zeiss), using a 63x/1.40 NA objective (Plan-Apochromat 63x/1.40 Oil DIC M27, Zeiss). Z stack images of GFP-positive neurons were acquired at a resolution of 17.45 pixels per µm in a z-step size of 0.29 µm. Identical pixel dwell time, laser power, pinhole size, and master gain were applied to all samples. Somata were segmented from the GFP channel using Otsu thresholding in ImageJ (version 1.54p), and the mean fluorescence intensity of PKA RIα within the soma was computed for each neuron.

### Quantification of PKA activity

Time-series images were registered using a SIFT feature-based alignment algorithm (Linear Stack Alignment with SIFT MultiChannel) in ImageJ. To segment the soma, a minimum-intensity projection of the 488 nm signal was generated, followed by Gaussian filtering (σ=1.0), and thresholding using “Auto Local Threshold” or “Auto Threshold” to produce optimal segmentation. Mean fluorescence intensity within the segmented soma was computed from the unfiltered 488 nm and 405 nm signals. Background fluorescence was estimated for each channel using five 50 x 50 pixel regions of interest (ROIs) devoid of cells. The mean background intensity was subtracted from the corresponding soma fluorescence intensity to obtain background-corrected fluorescence signals, *F_488_* and *F_405_*. Excitation-ratiometric PKA activity at each time point was calculated as the fluorescence ratio *R* = *F_488_*/*F_405_*. Changes in PKA activity were calculated as the normalized fluorescence ratio change, *ΔR*/*R_0_* = (*R* – *R_0_*)/*R_0_*, where *R* is the fluorescence ratio at a given time point, and *R_0_* is the fluorescence ratio at the first time point.

## Acknowledgements

We gratefully acknowledge Chan-Hyun Na, Taekyung Ryu, and David Kirchner at the Johns Hopkins Center for Proteomics Discovery for providing sample processing and LC-MS/MS services, and assistance with method description for this study. We express our heartfelt gratitude to Su-Jeong Kim and Solange Brown for their technical guidance for neuronal tracing, and to Aleksandr Smirnov and the Neuroscience Imaging Center in Johns Hopkins Department of Neuroscience for equipment and software support and guidance for neuronal tracing. This study was supported by the National Institutes of Health (R01MH129277 and U01DA056556), and the Maryland Stem Cell Research Fund.

## Contributions

AX Luo and PP Li conceived and designed the study. AX Luo and PP Li had full access to all data in the study and take responsibility for the integrity of the data and the accuracy of the data analysis. AX Luo performed biochemistry experiments, tissue collection, primary neuron experiments, and analyses of behavioral, biochemical, full-length RNA-seq, snRNA-seq, proteomic, PKA activity, and neuron morphometric data, and drafted the manuscript. L Deng conducted the mouse behavioral experiments. Y Li and J Li contributed to pilot behavioral testing, and Y Li, J Li, X Zhu, and CA Ross provided technical guidance for mouse behavioral testing. X Zhang contributed to pilot primary neuron experiments, and X Zhang and X Mao provided technical guidance for primary neuronal culture. B Wu provided technical guidance for PKA sensor design and imaging. CA Ross, B Wu, and Y Su reviewed the manuscript, and provided intellectual support and conceptual advice. PP Li supervised the study, provided funding support, and contributed to manuscript drafting. All authors have approved the final manuscript.

## Corresponding author

Correspondence to Pan P. Li.

## Ethics declarations

## Competing interests

The authors declare no competing interests.

---

**Supplementary Fig. 1.**
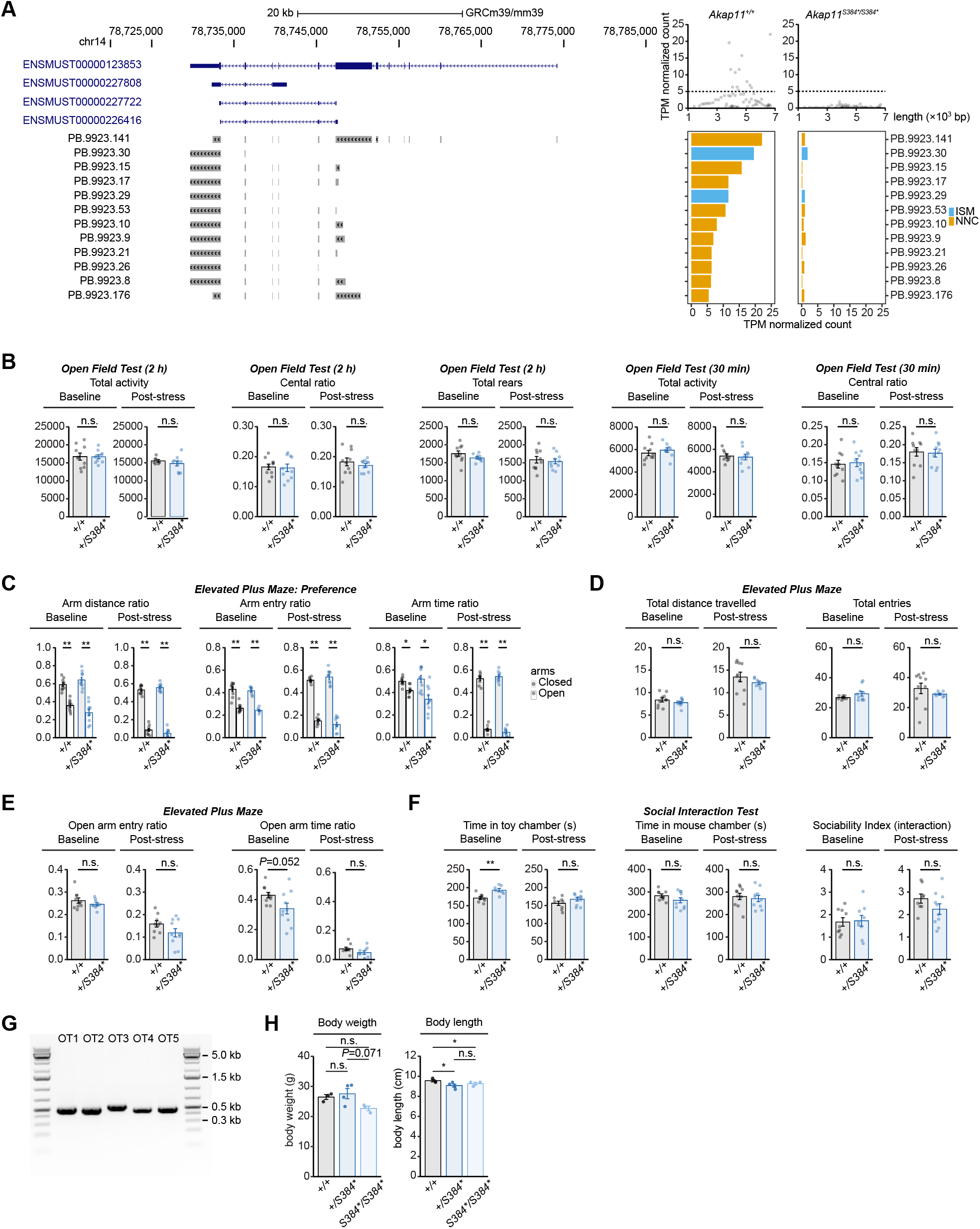
*Akap11* mRNA isoforms in homozygous mutant mice, and behavioral characterization of heterozygous mutant mice. Related to. Fig. 1. A. Genomic tracks showing four annotated *Akap11* transcript variants (blue) and *Akap11* isoforms with transcripts per million (TPM) ≥ 5 detected in the *Akap11^+/+^* cortical sample (gray) on the GRCm39/mm39 genome assembly. Upper right, *Akap11* isoform expression level, shown as TPM-normalized count, plotted against isoform length in *Akap11^+/+^* and *Akap11^+/S384*^* samples. Lower right, bar plots showing TPM-normalized expression levels for the isoforms displayed in the genomic tracks. ISM, incomplete splice matches, NNC, novel not in catalog. B. *Akap11^+/S384*^* mice showed normal total activity, central ratio, and total rears in 2-h open field test at baseline and after three weeks of isolated housing. No difference was detected between genotypes in total activity or central ratio during the first 30 min of the open field test. n.s. *P* > 0.05 via Mann-Whitney U test. Outlier removal and full statistical results are described in Supplementary Table 2A. C. Both *Akap11^+/+^* and *Akap11^+/S384*^* mice showed a preference for the closed arms in the elevated plus maze test, as indicated by the distance, entry and time measures. \*\**P* ≤ 0.01; \**P* ≤ 0.05 via paired Wilcoxon signed-rank test. Outlier removal and full statistical results are provided in Supplementary Table 2B. D. *Akap11^+/+^* and *Akap11^+/S384*^* mice did not differ in the total activity level in the elevated plus maze, measured by total distance travelled and total zone entries. n.s. *P* > 0.05 via Mann-Whitney U test. Outlier removal and full statistical results are provided in Supplementary Table 2B. E. *Akap11^+/S384*^* mice showed normal open-arm entry ratio in the elevated plus maze test at baseline and after isolated housing. Open-arm time ratio showed a trend toward reduction in *Akap11^+/S384*^* mice at baseline (*P* = 0.052) which was not observed after isolated housing. n.s. *P* > 0.05 via Mann-Whitney U test. Outlier removal and full statistical results are provided in Supplementary Table 2B. F. In the social interaction test, *Akap11^+/S384*^* mice spent significantly more time in the toy chamber than the control mice at baseline (*P* = 0.0081), but not after isolated housing. Time spent in the mouse chamber and interaction-based sociability index did not differ between genotypes before or after isolated housing. \*\**P* ≤ 0.01; n.s., *P* > 0.05 via Mann-Whitney U test. Outlier removal and full statistical results are provided in Supplementary Table 2C. G. Polymerase chain reaction (PCR) amplification products for the top five predicted sgRNA off-target sites (OT1-5). H. Bar plots showing effect of genotype on body weight (left) and body length (right) of 12-week-old male mice. Genotype had a marginal effect on body weight (*P* = 0.065 via a rank-transformed linear model with genotype and litter as fixed effects). *Post-hoc* analysis with Tukey’s test showed that homozygous mutant mice had marginally lower body weight than heterozygous mutant mice (*P* = 0.071), and no difference between other genotype pairs. Genotype significantly affected body length (*P* = 0.0035 via rank-transformed linear model with genotype and litter as fixed effects). Both heterozygous and homozygous mutants were shorter than control mice (*P* = 0.014 and *P* = 0.028, respectively, via Tukey’s test). No difference between the heterozygous and homozygous mutants was observed. \**P* ≤ 0.05; n.s. *P* > 0.05 via Tukey’s test.

**Supplementary Fig. 2.**
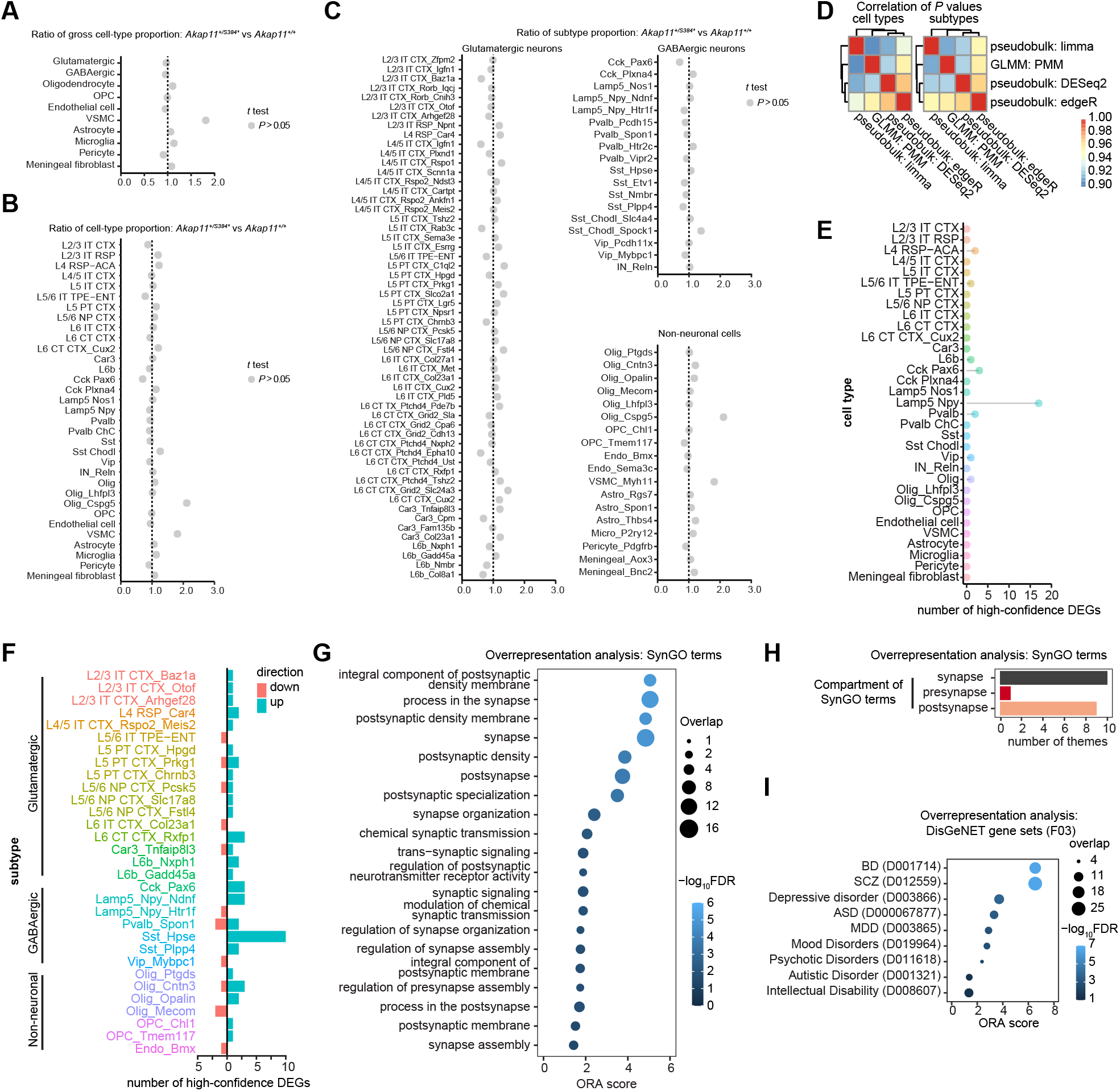
Single-nucleus RNA sequencing profiling of the neocortex of heterozygous mutant mice. Related to. Fig. 2. A. Ratio of broad cell type proportions between genotypes, calculated as *Akap11^+/S384*^* over *Akap11^+/+^*. Gray dots, *P* > 0.05 in cell type composition test via moderated *t*-test using *propeller*. B. Ratio of cell type proportions between genotypes, calculated as *Akap11^+/S384*^* over *Akap11^+/+^*. Gray dots, *P* > 0.05 in cell type composition test via moderated *t*-test using *propeller*. C. Ratio of subtype proportions between genotypes, calculated as *Akap11^+/S384*^* over *Akap11^+/+^*. Gray dots, *P* > 0.05 in cell type composition test via moderated *t*-test using *propeller*. D. Heatmap showing Pearson correlations among *P* values from three pseudobulk differential expression (DE) tests (via *edgeR*, *DESeq2*, and *limma*) and a single-cell Poisson-gamma mixed model (PMM)-based DE test (via *nebula*). Correlations are shown for the 33 cell types (left) and the 92 subtypes (right). E. Lollipop plot showing the number of high-confidence differentially expressed genes (DEGs) detected in each of the 33 cell types. High-confidence DEGs were defined as genes with FDR ≤ 0.10 in pseudobulk DE test via quasi-likelihood negative binomial generalized log-linear model with *edgeR*, and nominal *P* ≤ 0.05 in PMM-based single-cell DE test, with concordant direction of change between methods. F. Number of high-confidence DEGs detected in each subtype. Green indicates upregulated genes, and red indicates downregulated genes in *Akap11^+/S384*^* mice. Subtype labels are colored according to the cell type to which they belong. G. Synapse-specific gene ontology (SynGO) terms overrepresented among upregulated high-confidence DEGs across all subtypes. Significant overrepresentation was defined as FDR ≤ 0.05 via hypergeometric test. The overrepresentation analysis (ORA) score was calculated as −log_10_FDR multiplied by the direction of change, with positive values indicating enrichment among genes upregulated in *Akap11^+/S384*^* mice, and negative values indicating enrichment among downregulated genes. H. Distribution of overrepresented SynGO terms by synaptic compartment. I. DisGeNET gene sets in the MeSH F03 category (“mental disorders”) overrepresented among the extended high-confidence DEGs. Significant overrepresentation was defined as FDR ≤ 0.05 via hypergeometric test. The ORA score was calculated as −log_10_FDR multiplied by the direction of change. Abbreviations: ASD=autism spectrum disorder, BD=bipolar disorder, MDD=major depressive disorder, and SCZ=schizophrenia.

**Supplementary Fig. 3.**
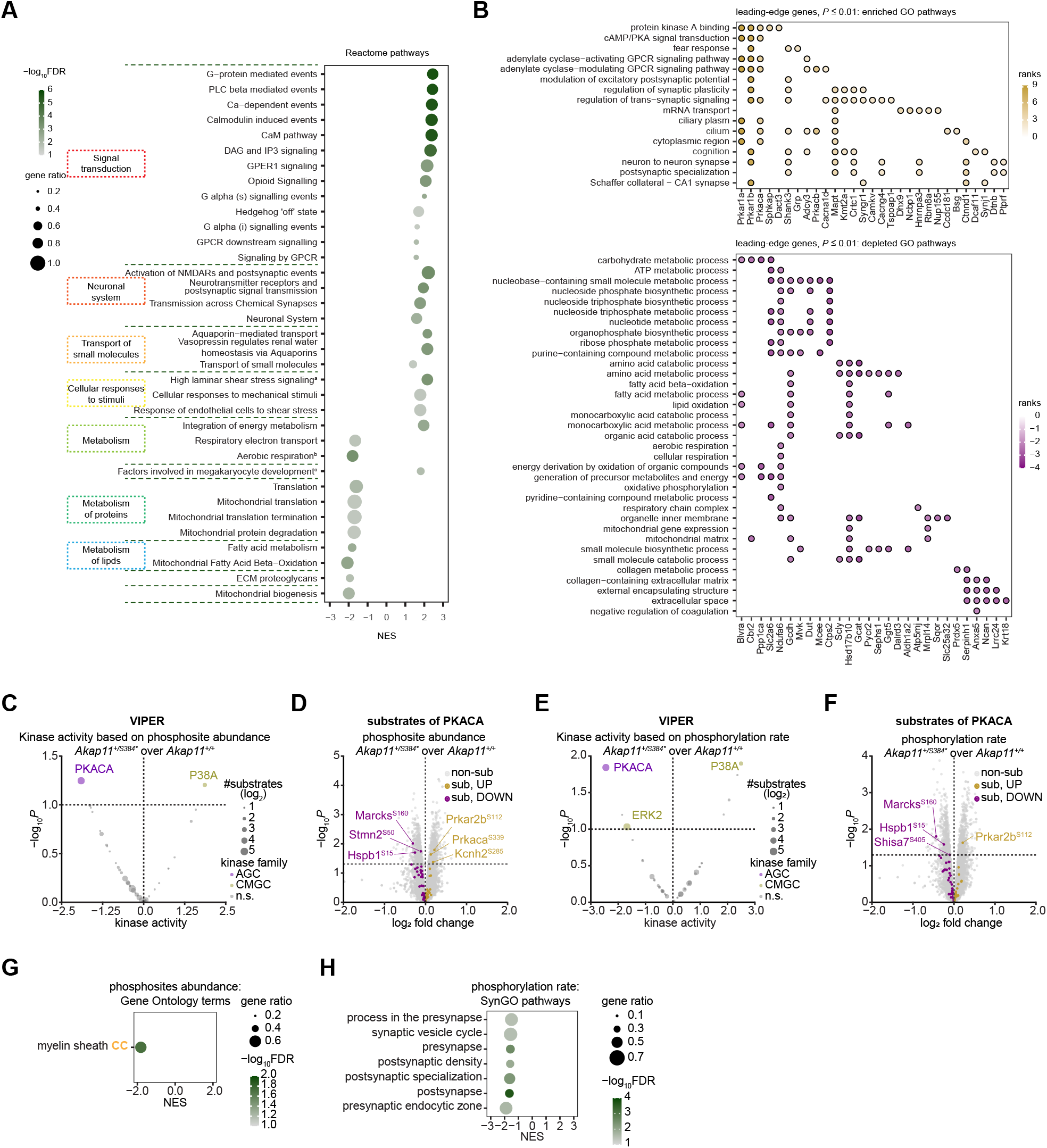
Proteomic profiling of the cortex of heterozygous mutant mice. Related to. Fig. 4. A. Reactome pathways significantly enriched or depleted in *Akap11^+/S384*^* mice via permutation test. Positive and negative normalized enrichment scores (NES) indicate enrichment and depletion, respectively in the heterozygous mice. Gene ratio was computed as the proportion of pathway genes represented in the ranked protein list. Significance was defined as FDR ≤ 0.05. Pathways are grouped into 10 groups according to the Reactome pathway hierarchy (retrieved from https://reactome.org/). Semantic summary are shown for categories containing multiple pathways. Truncated labels: ^a^High laminar flow shear stress activates signaling by PIEZO1 and PECAM1:CDH5:KDR in endothelial cells; ^b^Aerobic respiration and respiratory electron transport; ^c^Factors involved in megakaryocyte development and platelet production. B. Leading-edge proteins for the enriched (upper) and depleted (lower) Gene Ontology (GO) terms shown in Fig. 4C. Color indicates protein rank, which was computed as −log_10_*P* x sign (log_2_ fold change) from the differential protein abundance analysis. Positive and negative ranks indicate increased and decreased abundance, respectively, in *Akap11^+/S384*^* mice. Only leading-edge proteins with *P* ≤ 0.01 and GO terms with at least one such protein are shown. C, E. Volcano plots showing predicted kinase activity inferred from phosphosite abundance (C) or phosphorylation rate (E), using an alternative test method (VIPER). VIPER was applied to −log_10_*P* x sign(log_2_ fold change). Positive and negative kinase activity scores indicate increased and decreased kinase activity, respectively, in *Akap11^+/S384*^* mice. Kinases were considered significant at *P* ≤ 0.05 with at least three substrates present. D, F. Volcano plots showing genotype-dependent differences in phosphosite abundance (D), and phosphorylation rate (F), highlighting the predicted PKACA substrates associated with the *RoKAI* test. Positive and negative log_2_ fold changes indicate increased and decreased phosphosite abundance or phosphorylation rate, respectively, in *Akap11^+/S384*^* mice. Horizontal and vertical dashed lines indicate *P* at 0.05 and log_2_ fold change at 0, respectively. G. Gene set enrichment analysis (GSEA) of GO terms using phosphosites ranked −log_10_*P* x sign(log_2_ fold change), based on phosphosite abundance. Positive and negative NES values indicate pathways associated with proteins harboring increased or decreased phosphosite abundance, respectively, in *Akap11^+/S384*^* mice. *P* values were estimated via permutation test, and significance was defined as FDR ≤ 0.05. Gene ratio was computed as the proportion of pathway genes present in the phosphosite dataset. H. GSEA of SynGO terms using phosphosites ranked by −log_10_*P* x sign(log_2_ fold change) based on phosphorylation rate. Positive and negative NES values indicate SynGO pathways associated with proteins harboring increased or decreased phosphorylation rates, respectively, in *Akap11^+/S384*^* mice.

**Supplementary Fig. 4.**
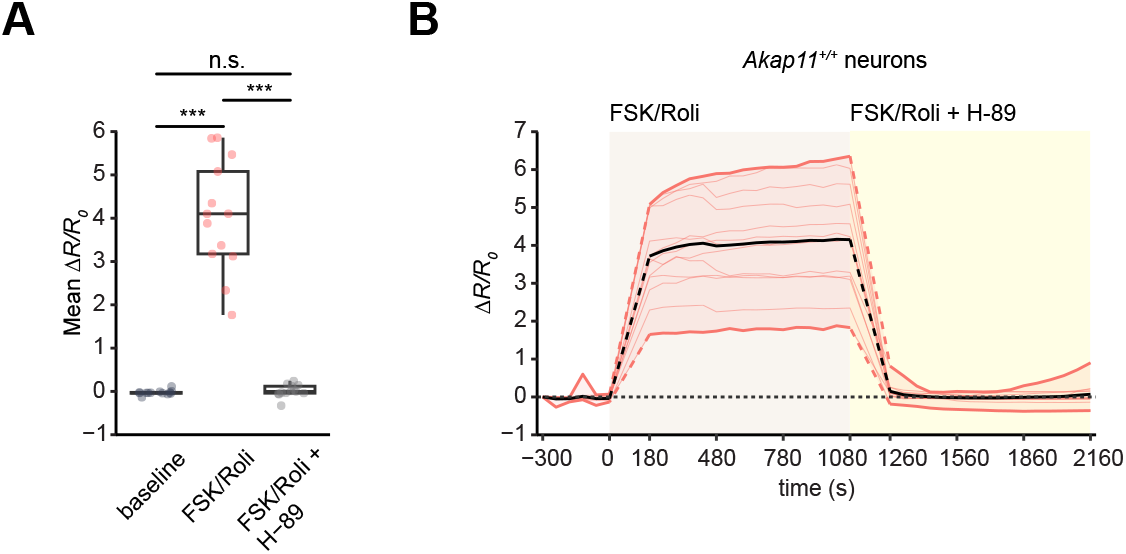
Validation of PKA sensor. Related to. Fig. 5. A. Boxplots showing the mean change in *F_488_/F_405_* ratio normalized to the first time point (mean *ΔR/R_0_*) in *Akap11^+/+^* neurons at baseline, in response to forskolin/rolipram (FSK/Roli), and in response to addition of H-89 in the continued presence of forskolin/rolipram (FSK/Roli+H-89). Forskolin/rolipram significantly increased sensor response relative to baseline (FDR = 7.32 x 10^-4^), whereas subsequent addition of H-89 restored the response to baseline (FSK/Roli+H-89 vs. FSK/Roli: FDR = 7.32 x 10^-4^; FSK/Roli+H-89 vs. baseline: FDR = 0.50). N = 2 embryos, n= 13 neurons. ***FDR ≤ 0.001, n.s., FDR > 0.05 via paired Wilcoxon signed-rank test. For boxplots, the bottom edge, midline, and top edge represent the first quantile, median, and third quantile, respectively. Whiskers represent 1.5 x interquartile range from the box edges. Full statistical results are shown in Supplementary Table 7B. B. Time course of change in *ΔR/R_0_* for the neurons shown in A. Dashed lines indicate intervals during drug application, when imaging was paused.

## Supplementary information

Supplementary Table 1

Supplementary Table 2

Supplementary Table 3

Supplementary Table 4

Supplementary Table 5

Supplementary Table 6

Supplementary Table 7

Supplementary Table 8

## Notes

### Competing Interest Statement

The authors have declared no competing interest.

### Summary of Updates

This version includes fixing typographical errors in Fig. 6, Supplementary Fig. 3, and Acknowledgements; minor rephrasing to reduce total word count to meet journal requirements; and addition of sample size and number of outliers removed in Fig. 1 legends.

